# Single-Cell Phenotyping of Extracellular Electron Transfer via Microdroplet Encapsulation

**DOI:** 10.1101/2024.06.13.598847

**Authors:** Gina Partipilo, Emily K. Bowman, Emma J. Palmer, Yang Gao, Rodney S. Ridley, Hal S. Alper, Benjamin K. Keitz

**Affiliations:** McKetta Department of Chemical Engineering, University of Texas at Austin, Austin, TX, 78712; Interdisciplinary Life Sciences Graduate Program, University of Texas at Austin, Austin, TX, 78712; Civil, Architectural, and Environmental Engineering, University of Texas at Austin, Austin, TX, 78712

## Abstract

Electroactive organisms contribute to metal cycling, pollutant removal, and other redox-driven environmental processes. Studying this phenomenon in high-throughput is challenging since extracellular reduction cannot easily be traced back to its cell of origin within a mixed population. Here, we describe the development of a microdroplet emulsion system to enrich EET-capable organisms. We validated our system using the model electroactive organism *S. oneidensis* and describe the tooling of a benchtop microfluidic system for oxygen-limited processes. We demonstrated enrichment of EET-capable phenotypes from a mixed wild-type and EET-knockout population. As a proof-of-concept application, bacteria were collected from iron sedimentation from Town Lake (Austin, TX) and subjected to microdroplet enrichment. We observed an increase in EET-capable organisms in the sorted population that was distinct when compared to a population enriched in a bulk culture more closely akin to traditional techniques for discovering EET-capable bacteria. Finally, two bacterial species, *C. sakazakii* and *V. fessus* not previously shown to be electroactive, were further cultured and characterized for their ability to reduce channel conductance in an organic electrochemical transistor (OECT) and to reduce soluble Fe(III). We characterized two bacterial species not previously shown to exhibit electrogenic behavior. Our results demonstrate the utility of a microdroplet emulsions for identifying putative EET-capable bacteria and how this technology can be leveraged in tandem with existing methods.

## Introduction

In the absence of oxygen, electroactive microorganisms export electrons out of the cell to reduce extracellular soluble and insoluble metal species in a process known as extracellular electron transfer (EET)^1^. This process is coupled to microbial growth, respiration, communication and sensing^2^. EET has been implicated in metal transport^3^, environmental remediation^4–7^, human health^8–11^ and more. Furthermore, EET has been co-opted for the treatment of wastewater^12,13^, power generation in microbial fuel cells^14–19^, and biocatalysis^20–24^. Electroactive bacteria have been isolated from a variety of locations including in aquatic and soil environments^12,25–27^, the human gut microbiome^8–10,28^, and oral biofilms^29^. Traditional methods for identifying EET-capable microbes involve competitive respiration on poised iron electrodes, or other metal-based electron acceptors^12,30–39^. Unfortunately, it is difficult to establish genotype-phenotype relationship, especially in complex consortia, because EET occurs in the extracellular space. Bulk enrichments using poised electrodes avoid this challenge by selecting for biofilms^16^ comprised of single or a handful of different electroactive species, but these experiments may not capture the complexity of the initial isolate^31,39–42^. A single species that forms a stable biofilm in bioelectrochemical cells can occlude other species from being incorporated into the biofilm altogether^43^. Such low throughput, bulk enrichment strategies require successful candidates outcompete other microbes for nutrients, electron acceptors, carbon sources, and access to the electrode. It also requires that a microbe be culturable in a laboratory environment. Consequently, bulk enrichment does not typically allow for more nuanced analysis of individual bacteria or the identification of EET in less fit, lower relative abundance, and unculturable microorganisms. Therefore, there is a need for high-throughput identification of single-cell EET behavior to identify novel electrogens in complex environments^32,44,45^.

Cytometric techniques, such as flow cytometry and flow-assisted cell sorting, are powerful techniques for characterizing complex populations but require intracellular reporters and cannot be easily translated to extracellular processes. Accordingly, microdroplet emulsions have recently emerged as a method for characterizing extracellular processes including cell surface display, secretion, and other processes^46^. Individual microbes can be statistically isolated in microdroplet systems by diluting them in an aqueous solution and forming microdroplet emulsions via an oil countercurrent flow. Once formed, aqueous droplets can be cultured, merged with fresh aqueous solutions, and analyzed via fluorescence where they can be sorted into populations of low and high performers. Essentially, single-cell aqueous microdroplets function as individual reaction wells, capturing extracellular products or fluorescence^47^ and muting cell-to-cell competition. Microdroplet screening of microbial consortia has previously been used for bioprospecting phenotypes of interest from environmental populations^48–50^, but to the best of our knowledge has not been used to identify electroactive organisms. This is likely due to the additional challenge of maintaining anaerobic conditions within a microdroplet screen and the lack of fluorescent reporters for EET. However, recently we detailed the coupling of Cu(I)-catalyzed Alkyne-Azide Cycloaddition (CuAAC) to electron transfer from *S. oneidensis*^24^, and saw that fluorescence from the cycloaddition of a small molecule could be used to quantify electron transfer.

Here, we describe the development of a high-throughput microdroplet assay for characterizing EET. We detail the adaptation of CuAAC as a mechanism for EET-detection and the tooling of microdroplet instrumentation for use with anaerobic cultures via an oxygen-limited protocol. Our system was developed utilizing a monoculture of model electroactive organism *Shewanella oneidensis* MR-1, which allowed us to assess assay performance. A mixed population of EET-deficient strains and EET-capable strains of *S. oneidensis* facilitated fluorescent activated droplet sorting (FADs) to enrich the EET-capable strain from the mixed population. Additionally, a defined population mixture of *E. coli* Nissle 1917, *S. cerevisiae* BY4741, and *S. oneidensis* MR-1 demonstrated that signal could be detected even from a complex sample. As a test application, we identified EET-capable microbes from an environmental sample. We saw a distinctly enriched EET-capable population, which had upregulation of iron reduction genes compared to the initial consortium. Two bacteria, *Cronobacter sakazakii* and *Vagococcus fessus*, identified by our microdroplet screened were assessed in monoculture, and displayed marked electrogenic behavior. Herein, we describe the development of a microfluidic method for identifying extracellular reduction by EET-capable bacteria that is does not rely on reduction coupled to competitive growth.

## Results

### Optimization of CuAAC fluorescent probe assay for applications in microdroplets

We recently utilized fluorescence from the EET-driven synthesis of a cycloaddition probe, CalFluor488^51^, to assess EET activity from the bacteria *S. oneidensis* via Cu(I)-catalyzed alkyne-azide cycloaddition (CuAAC)^52^. We determined that the rate of copper reduction was directly correlated to EET flux via the well-defined EET-protein pathway in *S. oneidensis* (the Mtr-pathway), and that fluorescence was tied to electron transfer. As a result, we recognized that the system could be used as a fluorescent sensor for EET, and we hypothesized that the fluorescence readout could be adapted into a screen for identifying electroactive microbes in microfluidic droplets. The quenched to fluorescent transition upon cycloaddition of the CalFluor488^51^ probe is ideal for microfluidic application as the unreacted emulsion has little background fluorescence (Figure 1, Figure S2). First, we confirmed that neither the reaction nor the fluorescence output was inhibited by the reagents required for a stable emulsion. Traditionally, both a rich media and low-to-no organic solvents are required for stabilizing emulsions as well as supporting microbial growth within droplet emulsions^46^. Our previous use of *S. oneidensis* to perform CuAAC was performed in minimal media with DMSO as a co-solvent^52^, and which in initial screens yielded droplet instability. Anaerobic reactions performed with *S. oneidensis* in a 96-well plate showed that nearly all DMSO could be omitted apart from the small volume of co-solvent in the CalFluor488 stock. Neither cell growth nor conversion was hindered under these conditions even with increased concentrations of reagents. (Figure S1). A similar control performed in Lysogeny broth (LB broth) in the presence of fluorinated oil with or without the biosurfactant present (Sphere fluidics), showed a slightly decreased, but still large fluorescence response.

**Figure 1.**
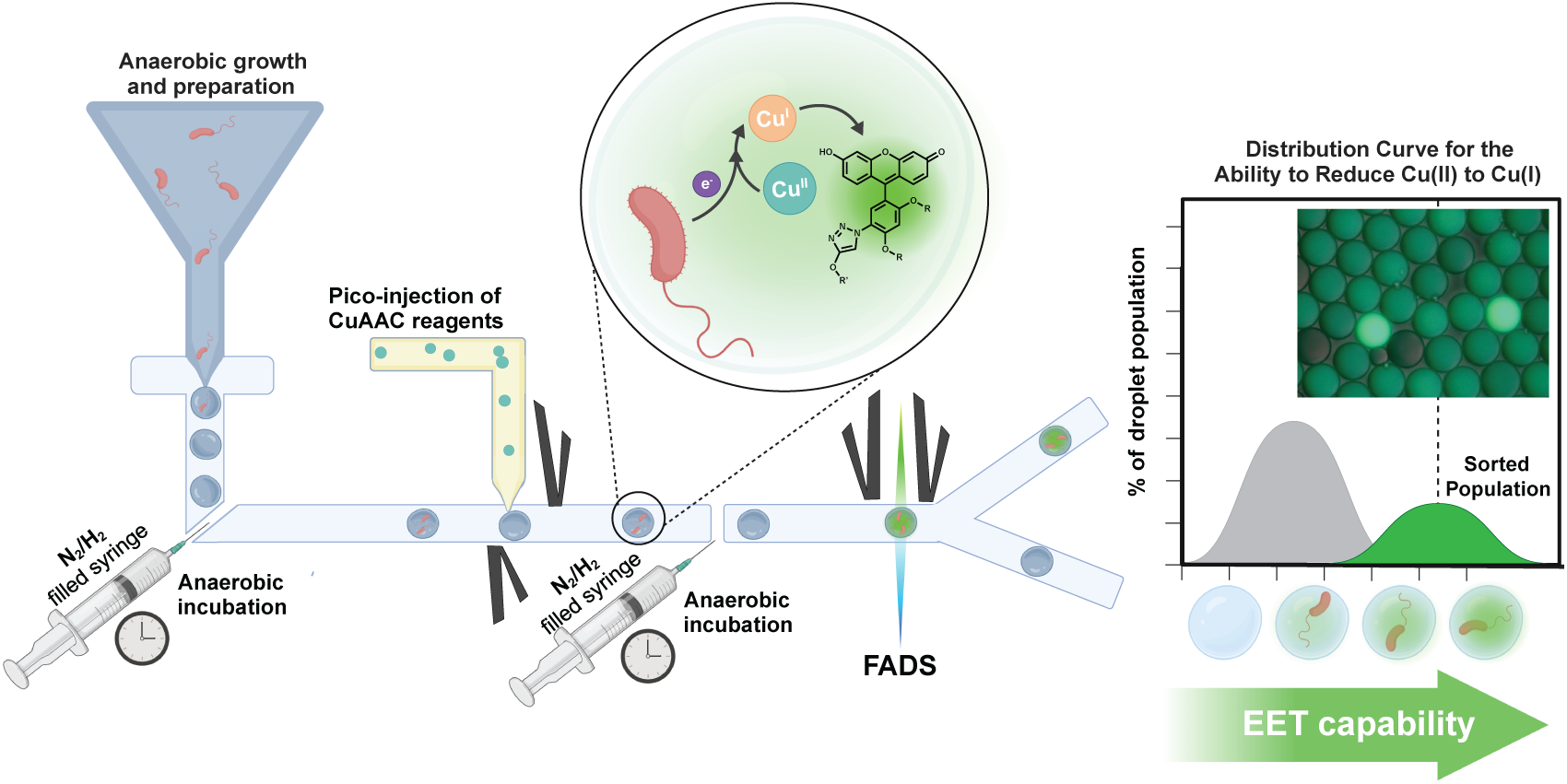
Microdroplet assay set up and schematic. The reaction scheme for performing oxygen-limited CuAAC via microbial respiration in a microfluidic system. Inset is an image of *S. oneidensis*-containing microdroplets image prior to sorting.

Having confirmed that the reagents necessary for a stable emulsion did not interfere with cell growth or CuAAC activity, we moved forward to adapting the assay to the microdroplet system. Abiotic emulsions were created with starting material and chemically-synthesized product. The emulsions were mixed in known ratios and analyzed via fluorescent activated droplet sorting (FADS) to determine the ability to distinguish reacted product from background (Figure S2). We found that the system could detect differences in the population including the 10% reacted emulsion from the 90% unreacted emulsion which is the target level of cellular encapsulation during a screen (1 filled droplet in every 10). Together, these benchtop assays suggested the CuAAC chemical system could be utilized as a microbially-driven fluorescent readout for microdroplet emulsion sorting.

### Oxygen-limited benchtop microdroplet system enabled Cu(I)-catalyzed Alkyne-Azide

Wild-type *S. oneidensis* can convert CalFluor488, to a fluorescent cycloaddition product in approximately 5 hours when combined and sealed with the appropriate Cu, ligand, and alkyne source^52^. We began by emulsifying an aqueous reaction containing aerobically grown *S. oneidensis* at an OD_600_ of 6 x 10^-5^, calculated such that one in every 10 droplets was filled^53^, using the fluorinated oil and biosurfactant as the oil phase. However, we measured no distinguishable fluorescent signal after 24 hours under these conditions. Hypothesizing that the dilute cells could not withstand the simultaneous stress of encapsulation in the presence of Cu(II/I) and small molecules, we adapted a previously developed method for utilizing biosensors in microdroplets^46^. Specifically, we utilized “pico-injection” which flows a previously formed emulsion through a microfluidic chip and introduces a fresh aqueous solution into each droplet by applying a low voltage to merge the emulsion with the aqueous pico-injection solution. Previously, microdroplet emulsions have been pico-injected to add cell-based biosensors^46^, cell lysis reagents^54,55^, or fluorogenic enzyme substrates^56–59^ into existing emulsions. We hypothesized this mechanism would provide robustness for withstanding the stress of both the microdroplet system and the Cu(II/I) and small molecules (Figure S3). We moved to pico-inject the CalFluor488, Cu(II), ligand, and alkyne after the emulsion had been created and allowed to stabilize for a full day. The 5X concentrated pico-injection solution was flowed into the emulsion utilizing a Pico-Mix^TM^ chip (Sphere Fluidics) and merged under a low voltage of 0.15V where it was diluted to the desired concentration. This facilitated growth within the droplets, where the cell concentration within the emulsion increased, but each droplet contained genetically identical cells^56^ with no droplet-to-droplet competition for resources. However, we measured no detectable fluorescent signal even under pico-injection scheme (Figure 2a). We posited that the emulsion was too oxygen-permeable, and that oxygen was preventing a shift to anaerobic metabolism in *S. oneidensis* as well as inhibiting the CuAAC reaction.

**Figure 2.**
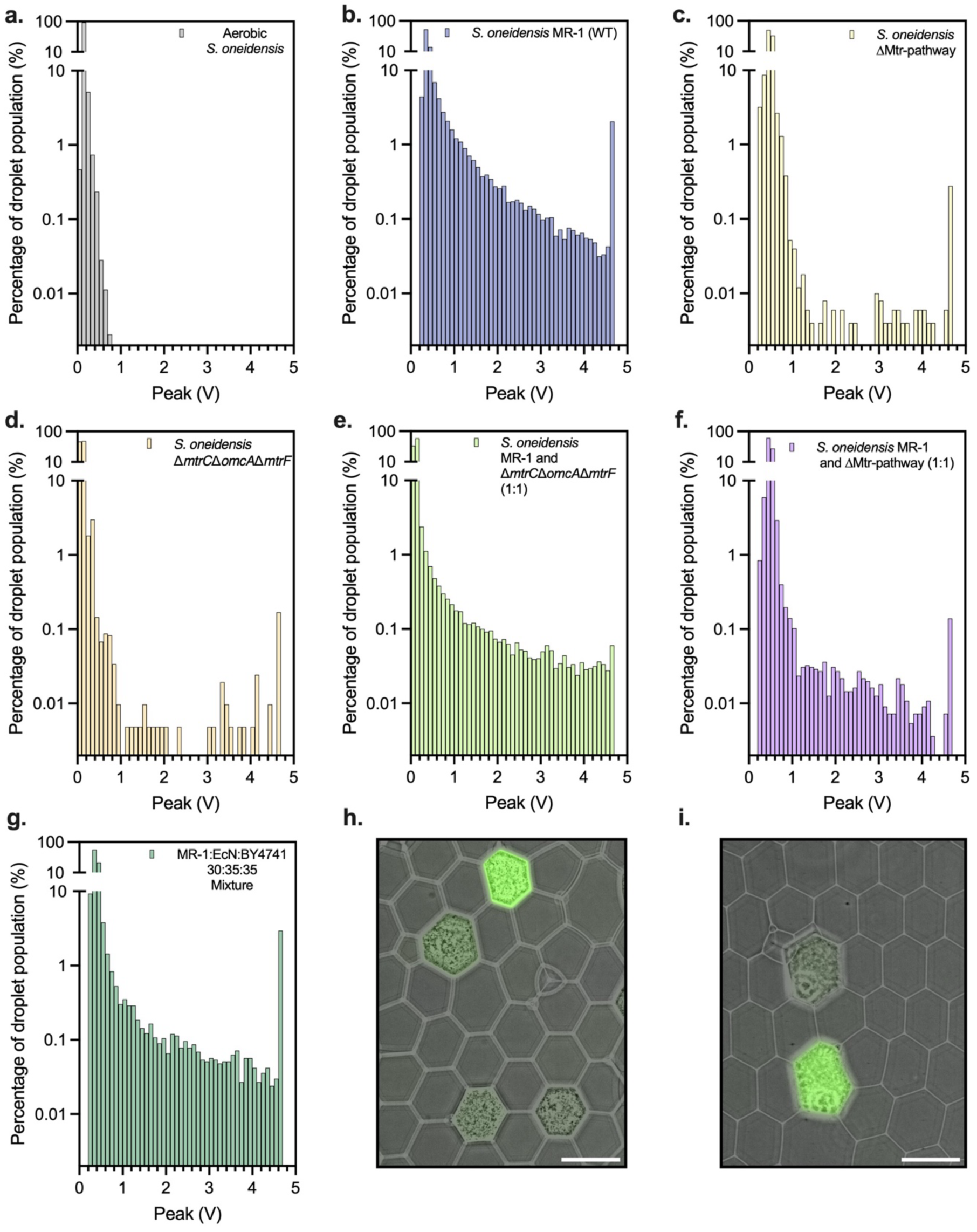
Oxygen-limited conditions allow for detection using of *S. oneidensis* in microdroplet system. **a.** Histogram of aerobic FADS of *S. oneidensis* **b.** Histogram of an oxygen-limited FADS of *S. oneidensis*. **c.** Histogram of an oxygen-limited FADS of ΔMtr-pathway *S. oneidensis* **d.** Histogram of an oxygen-limited FADS of Δ*mtrC*Δ*omcA*Δ*mtrF S. oneidensis.* **e.** Histogram of an oxygen-limited FADS of *S. oneidensis* wild-type and *mtrC*Δ*omcA*Δ*mtrF S. oneidensis* mixed in a 1:1 ratio prior to encapsulation. **f.** Histogram of an oxygen-limited FADS of *S. oneidensis* wild-type and *S. oneidensis* ΔMtr-pathway mixed in a 1:1 ratio prior to encapsulation. **g.** Histogram of an oxygen-limited FADS of *S. oneidensis* wild-type, *S. cerevisiae* BY4741, and *E. coli* Nissle 1917 mixed in a 30:35:35 ratio prior to encapsulation. **h.** Representative image of emulsion from (**g.)** prior to sorting with merged bright-field and fluorescence. **i.** Representative image of emulsion from (**f.)** prior to sorting with merged bright-field and fluorescence.

To overcome this challenge, we performed subsequent experiments in an oxygen-limited environment. All oils and buffers were sparged and prepared anaerobically. Tubing was attached, and the syringes were sealed within an anaerobic chamber to create a closed environment. A 12- mL collection syringe was similarly prepared with a small, cushion of fluorinated oil (500 µL) and anaerobic atmosphere (6 mL of 97% N_2_, 3% H_2_), then sealed with a needle and heat-sealed tubing. The solutions and oils were removed from the chamber and loaded onto the syringe pumps under a positive pressure (Figure S3). Once the system had equilibrated and the emulsion or pico- injection was stable in size, the collection syringe could be used to collect the emulsion. As predicted, this oxygen-limited set up allowed *S. oneidensis* MR-1 encapsulated within droplets to perform CuAAC, which could be detected using our system (Figure 1, Figure 2b). The fluorescence in the emulsion was clearly visible under fluorescent microscopy (Figure, 1 inset) and the ratio of fluorescent to non-fluorescent droplets approximated the predicted encapsulation ratio, one out of every 10 droplets filled. The histogram demonstrated that a higher percentage of the droplet population (y-axis) exhibited fluorescence (x-axis) and indicated we were able to detect the fluorescence on the FADS system (Figure 2a and 2b). In total, these results indicate that the bacteria can performs sufficient EET within the microdroplet system to catalyze CuAAC and that we can detect the extracellular reduction via fluorescence.

### CuAAC for the detection of Extracellular Electron Transfer in microdroplets

Next, to confirm that signal was tied to EET, an EET-deficient strain of *S. oneidensis* (ΔMtr- pathway) lacking all outer membrane cytochromes was subjected to droplet encapsulation and fluorescent screening. As expected, no significant fluorescence was detected (Figure 2c). To confirm the cells were still alive within the droplets, a voltage was applied, and the emulsion was broken to harvest the cells. After plating the cells onto agar plates, lawns were recovered for the sorted indicating the cells were alive within the droplets, despite their lack of signal (Figure S4). Furthermore, a knockout of only the terminal EET-proteins (MtrC, MtrF, and OmcA) resulted in a similar histogram without appreciable signal (Figure 2d). Having demonstrated that CuAAC within microdroplets could distinguish between homogeneous populations in individual emulsions, we next examined a mixed culture of wild-type and EET-deficient strains of *S. oneidensis* (Figure 2e and 2f).

A 1:1 ratio of wild-type MR-1 to knockout was mixed immediately prior to encapsulation. Immediately prior to sorting, microscopy revealed that approximately one out of every two filled droplets fluoresced (Figure 2i), indicating that that ΔMtr-pathway strain could be differentiated from wild-type *S. oneidensis* by fluorescence. During FADS of the mixture of Δ*mtrC*Δ*mtrF*Δ*omcA* and wild-type *S. oneidensis*, a subsection of the high fluorescent population was collected, and the emulsion was broken and plated on agar plates. Sections of the sorted emulsion were subjected to a colorimetric Fe(III/II) reduction assay^20^, and after 24 hours 9 colonies from each section were analyzed to determine if they had reduced Fe(III) to Fe(II). The unsorted population yielded 4 out of 9 colonies within one standard deviation of wild-type MR-1, while the sorted population (1-2V) had 7 colonies of MR-1, an enrichment of 1.75-fold over the starting material (Figure S5). These results validated that the microdroplet system could enrich EET-active strains in a mixed population. Wild-type could be enriched via fluorescence despite the fact that the Δ*mtrC*Δ*omcA*Δ*mtrF* strain is known to have spurious background reduction, due to the presence of the MtrA and MtrD which can still interact with soluble metals^22,52^. Based on the success at differentiating between two strains of the same species, we investigated the ability to detect an EET-capable bacteria within a synthetic multi-species co-culture. In picking our proof-of-concept consortia, we chose *Eschericia coli* Nissle 1917, *Saccharomyces cerevisiae* BY4741, and *S. oneidensis* MR-1 for their availability and ability to be grown at 30 °C, anaerobically. Each microbe was grown separately overnight, and mixed in a known ratio (35%, 35%, 30% respectively) immediately prior to encapsulation. We detected fluorescence even from this mixed population (Figure 2g) suggesting that our system could be used to characterize more complex samples. Under microscopy, visible growth was detected in approximately 10% of the microdroplets, which was the target encapsulation percentage. Of those with visible growth, approximately one in every three fluoresced (Figure 2h), indicating that even in more complex consortia, there is detectable differences within a population. These data indicate that emulsions made from homogenous populations are distinctly different and simple model consortia of very similar cells can be fluorescently sorted to isolate EET-capable phenotypes.

### Single-cell analysis of environmental samples reveals EET-capable bacteria

Next, to evaluate the microdroplet system for a more complex environmental sample and to compare to a more traditional screen for EET, bulk enrichment, we interrogated a sediment-associated mixed microbial community collected from Town Lake in Austin, TX. To compare the efficacy of our microdroplet method relative to alternative methods for screening for EET activity, part of the sample was anaerobically incubated with a solid-phase Fe(III) source to select for electroactive, EET-capable microbes. The bulk enrichment experiment included lactate as a carbon source and iron-rich sediment as the electron acceptor. Sterilized sediments were isolated within a 3.5 kDa MWCO dialysis membrane tube to limit the contact between bacteria and iron minerals and avoiding biofilm formation^58,60–62^. The bulk enrichment culture was monitored for both pH and soluble Fe(II) concentration overtime (Figure 3b) and displayed an increase in soluble Fe(II) from 0.069 mg/L to 0.8 mg/mL over the course of 5 days indicating that this biological sample contained EET-capable organisms. Samples from before and after enrichment were harvested and subjected to 16S sequencing (Figure 3, samples I and II).

**Figure 3.**
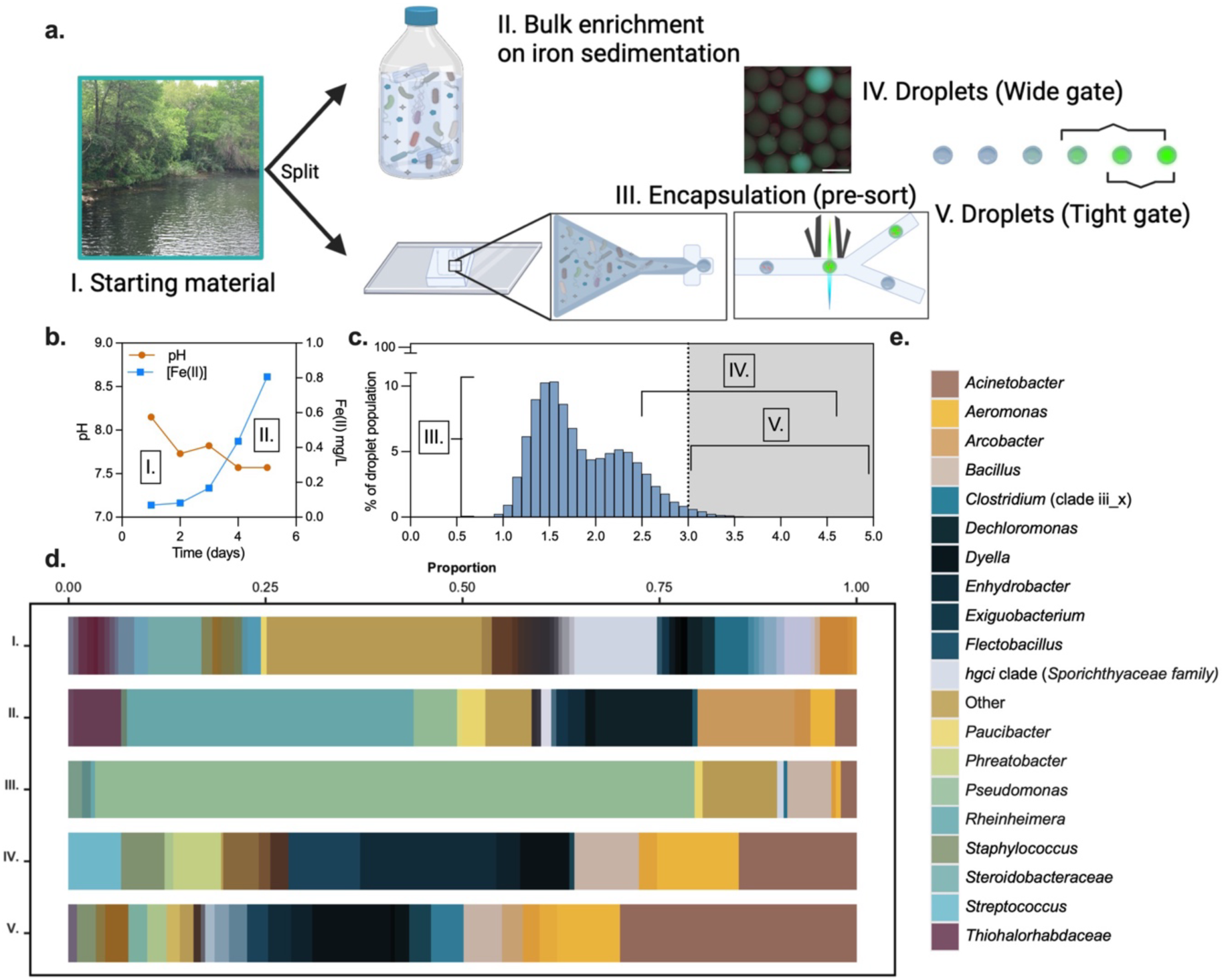
Iron sediment gathered from Town Lake yields Fe(III)-reducing bacteria. **a.** A schematic outlining the experimental procedure, and where samples were gathered for 16S sequencing. Briefly, bacteria were collected from iron sedimentation. This starting material was sequenced and split into a bulk enrichment and a microdroplet enrichment. The microdroplet samples was sequenced pre- and post-sorting. Two different gates of the sorted enrichment were collected and subjected to sequencing. Inset of droplets is a representative image of the lake water emulsion prior to sorting and scale bar represents 100 µm. **b.** Bulk enrichment data monitoring pH and Fe(II) concentration over time. **c**. Histogram of microdroplet emulsion and gating of samples IV. and V. Histogram represents relative proportion of population after system has stabilized and collected for 87k droplets. Sorted populations were collected for a minimum of 500 droplets collected. **d.** Relative genus distributions of samples with greater than 0.05% prevalence. **e.** Top 20 genus’s as describe in **d.**

### 16S sequencing allowed for identification of high-performing targets

The other section of split sediment-derived sample was subjected to the microfluidic screen where a clear sub-population exhibited improved fluorescence (Figure 3a), as indicated by a tailing end of high performers (Figure 3c). To evaluate the effect of microdroplet formation, both the initial sediment-derived population (Sample I) and the population immediately prior to sorting (Sample III) were both collected for 16S sequencing. To evaluate the effect of sorting the population by fluorescence, two portions of the population were sorted out, wide (greater than 2.5 V) (Sample IV) and tight (greater than 3.1V) (Sample V), gated and were also collected for 16S sequencing.

The 16S sequencing data was normalized by the number of reads per sample, and compared to the starting, untreated lake water. From the bulk enrichment (Sample II), 68 bacteria were identified and 20 (29.4%) of these had been previously thought to been capable of EET^26,31,37,44,63–66^. The remaining 48 could indicate “cheaters”, bacteria that survived in bulk but were not actively reducing iron upon harvest. The bulk enrichment favored the *Rheinheimera* and *Dechloromonas* and *Arcobacter* genera. In the microdroplet system, when compared to the initial lake water, 45 species were enriched by encapsulation alone. A significant portion of the population that was present in the unsorted microdroplets screen were members of the *Psuedamonas* genus. However, the sorted droplet populations were enriched and favored *Exiguobacterium*, *Acinetobacter*, and *Aeromonas* genera, indicating that the sorting selects for reduction as opposed to selecting exclusively for survival within droplets. We identified a total of 56 bacterial species that were identified in the fluorescently sorted droplets indicative of an ability to reduce Cu(II) to Cu(I). Of the 56 bacterial species identified in the droplet sorted population, 16 (28.6%) had been previously reported as potentially EET-capable^26,31,37,44,63–66^. However, of the 366 species bacteria detected in the starting material (or in a subsequent enrichment), only 49 (13.4% of the starting material) were putative EET-capable bacteria and, only 29 (57.1% of the previously reported) of those survived encapsulation prior to sorting. This indicated that some were either unable to withstand the sediment to microdroplet transition or did not survive the initial reconstitution from iron sedimentation. Given that only 29 putative EET-capable species survived encapsulation, 16 (55.2%) of the putative EET bacteria within the pre-sort droplets were identified by our screen indicating that the screen was effective for detecting EET activity (Figure 4b).

**Figure 4.**
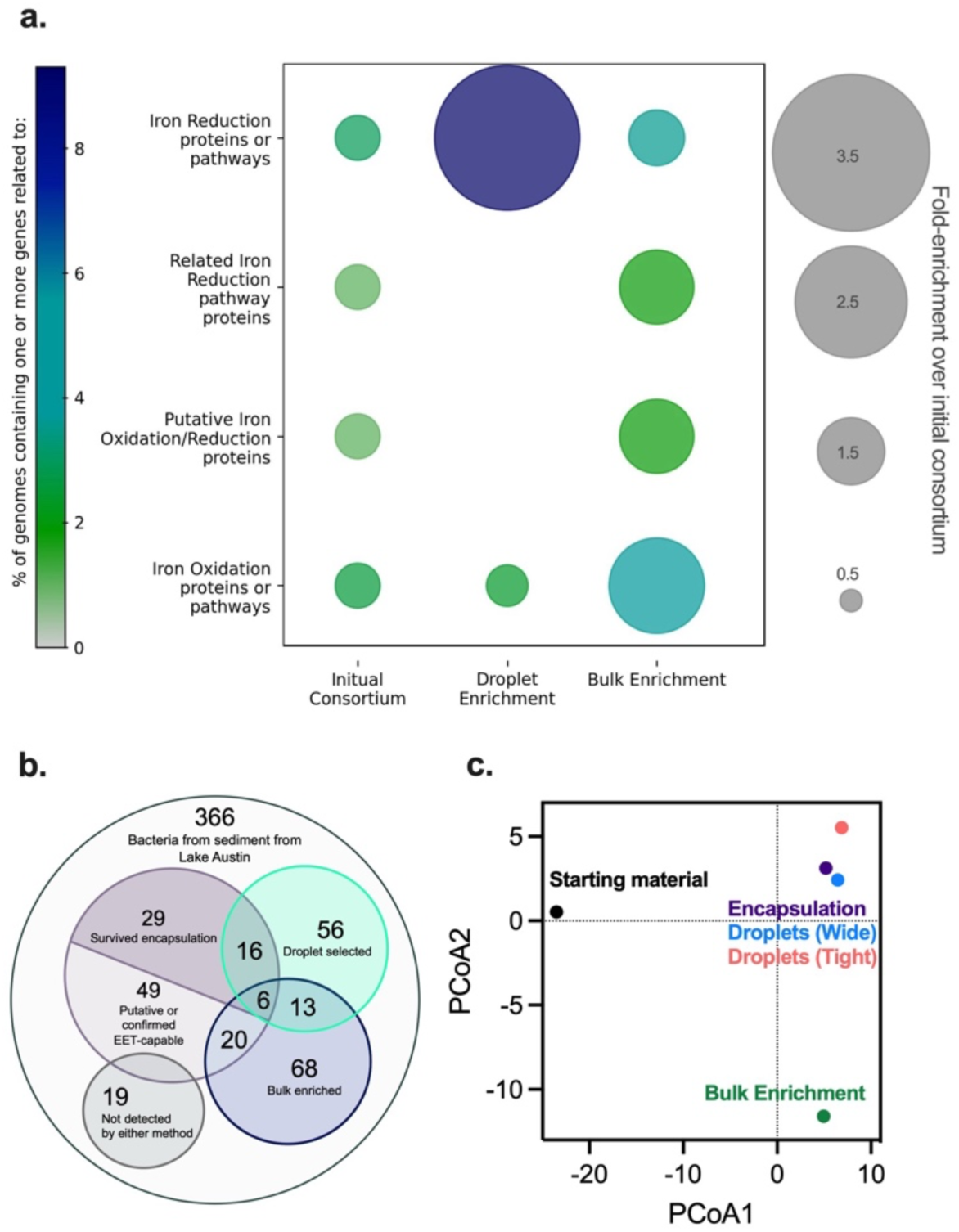
Analysis of species similarity between enrichment mechanisms. **a.** Percent of species containing one or more genes related to iron-regulation pathways as determined via HMM with FeGenie^67^. **b.** Venn diagram describing the relative species distributions and overlap between samples. **c**. Principal component analysis of the genus data represented in Figure 3d with a *k*=2.

Interestingly, when comparing between the microdroplet enrichment (Samples IV and V) and bulk enrichment (Sample II) only 13 bacteria were enriched under both conditions. Only 6 of these shared species had been previously referenced as EET-capable: *Acinobacter johnsonii, Aeromonas salmonicida, Aeromonas veronii*, *Ralstonia solanacearum,* an unidentified *Aeromonas* species and a *Citrobacter* species, and were identified by both droplet and bulk screens. 7 other species were identified by both methods but had never been characterized as having the ability to perform EET (Figure 4b). A principal component analysis (*k*=2) revealed that the starting material, droplets, and the bulk enrichment were clustered separately (Figure 4c), indicating that the populations of these samples were distinctly different. Finally, Rarefaction curves (Figure S6) suggested a drop in diversity between the starting material and the encapsulation, as well as the encapsulation and the sorted samples indicating that there was a bottleneck that decreased sample diversity. Together, this 16S sequencing data indicates that we enriched for a distinct sub-section of the population in the microdroplet FADS that potentially represents a phenotype with the ability to perform EET.

### FeGenie analysis of sorted populations reveal differing enrichments

To further probe the features of the populations identified by the bulk and the microdroplet enrichments, example genomes from each enriched bacteria (for which a fully sequenced genome was available) were analyzed using FeGenie to identify iron-trafficking genes.^67^ This program looks for homology using a hidden Markov model (HMM) against known iron-related protein pathways. We used it to identify whether a given bacterium within a population has one or more genes related to a known iron-redox pathway. Looking at four of the gene clusters related to iron reduction, the microdroplet system enriched for bacteria containing known pathways involved in iron reduction (CymA, MtrCAB, OmcF, OmcS, OmcZ, FmnA-dmkA-fmnB-pplA-ndh2-eetAB-dmkB^44,68–71)^, notably including those most similar to what is found in *S. oneidensis*^1,72–75^. Compared to the initial consortium, 3.23-times more bacteria had one or more genes related to this specific iron reduction pathway, nearly 10% of the genomes analyzed (Figure 4a). The program also screens for genes related to iron reduction pathways where the there is no homolog to the terminal *mtrC*. These are characterized as “related to iron reduction” because they represent a potential, but incomplete pathway^67–71^. None of the genomes analyzed from the droplet enrichment fell into this category; however, these bacteria containing “related to iron reduction” genes (MtrCB, MtrAB, MtoAB-MtrC) were enriched over 1.65-fold over the initial consortia in the bulk enrichment. These data probe into the possible molecular mechanism of enrichment, indicating that a terminal protein for metal-interaction may potentially be required in the microdroplet system. However, given that the mechanisms of EET are diverse, these data only capture genes prevalent in previously characterized electrogens. Interestingly, in the bulk enrichment, we measured an increase in presence of genomes containing one or more genes related to iron oxidation. We believe this was likely due to reduction of Fe(III) to Fe(II) in the bulk enrichment, which could be used for iron oxidation by other bacteria. Combined, these data suggest competing mechanisms of enrichment that likely may contribute to the low number of shared bacteria between the enrichment samples (Figure 4a). Additionally, this indicates that the microdroplet enrichment was uniquely distinct, and potentially represented a more specific population of electrogens compared to the bulk enrichment which may select for interspecies collaborations or sub-sets of EET-mechanisms that represent multiple simultaneous phenotypes.

### Monoculture characterization of high-performers from microdroplet enrichment reveal putative electrogens

Finally, we determined whether select species isolated in the highest performing droplets sort were EET-capable or if we were enriching for alternative phenotypes. To examine this, two bacteria identified solely in the highest sort (tight sort, Sample IV) gating of the lake water microdroplet enrichment, *Cronobacter sakazakii* and *Vagococcus fessus*, were characterized for their ability to perform EET. Neither bacteria was enriched in the bulk enrichment nor had they previously been reported as EET-capable, although *C. sakazakii* has been known to have iron-transport and acquisition related machinery that is vital to its survival^76,77^. Each bacterium was grown in its’ preferred culture media. As a benchmark, *E. coli* MG1655 and *S. oneidensis* MR-1 were measured concurrently in each respective media and evaluated for their EET-activity. The bacteria were incubated in an organic electrochemical transistor (OECT) under continuous electrode bias conditions to examine their ability to reduce insoluble electron acceptors. OECTs can translate and amplify biological signals into electrical responses where direct or shuttling EET reduces the conductance of the p-type channel^78^. OECTs have a faster response and require smaller volume compared to traditional electrochemical cells^78^. Conductance was measured under constant bias voltages (V_DS_ = -0.05V, V_GS_ = 0.2V). Both *C. sakazakii* and *V. fessus* exhibited a marked drop in conductance over the course of a 24 h period; however, only *V. fessus* outperformed *E. coli* in these devices (Figure 5a and 5b). To obtain a precise measurement of the electroactive activities, the transfer curves were plotted against the Ag/AgCl reference electrode (RE) (Figure S8). A more positive effective gate voltage (V_G_^EFF^) or a more negative measured source electrode potential (V_S_) indicates a reduction after incubation with the bacteria^78^. Next, to determine their ability to reduce soluble Fe(III), a Fe(III/II) reduction assay and growth on Fe(III) was collected for each bacteria. Both bacteria reduced Fe(III) to Fe(II) over 20 hours as measured by the colorimetric ferrozine assay^79^ (Figure 5c and 5d). Neither bacteria appeared to require Fe(III) strictly as an electron acceptor for anaerobic growth; however, we were unable to establish culturing conditions for *V. fessus* without the presence of at least 0.1% sheep’s blood (Figure 5e and 5f, Figure S7). In tandem these results suggest that these bacteria are capable of reducing soluble Fe(III) to Fe(II) and that, under our conditions, *V. fessus* exhibits similar EET-levels to *S. oneidensis* in reducing insoluble electron acceptors at the gate and channel of the OECT devices.

**Figure 5.**
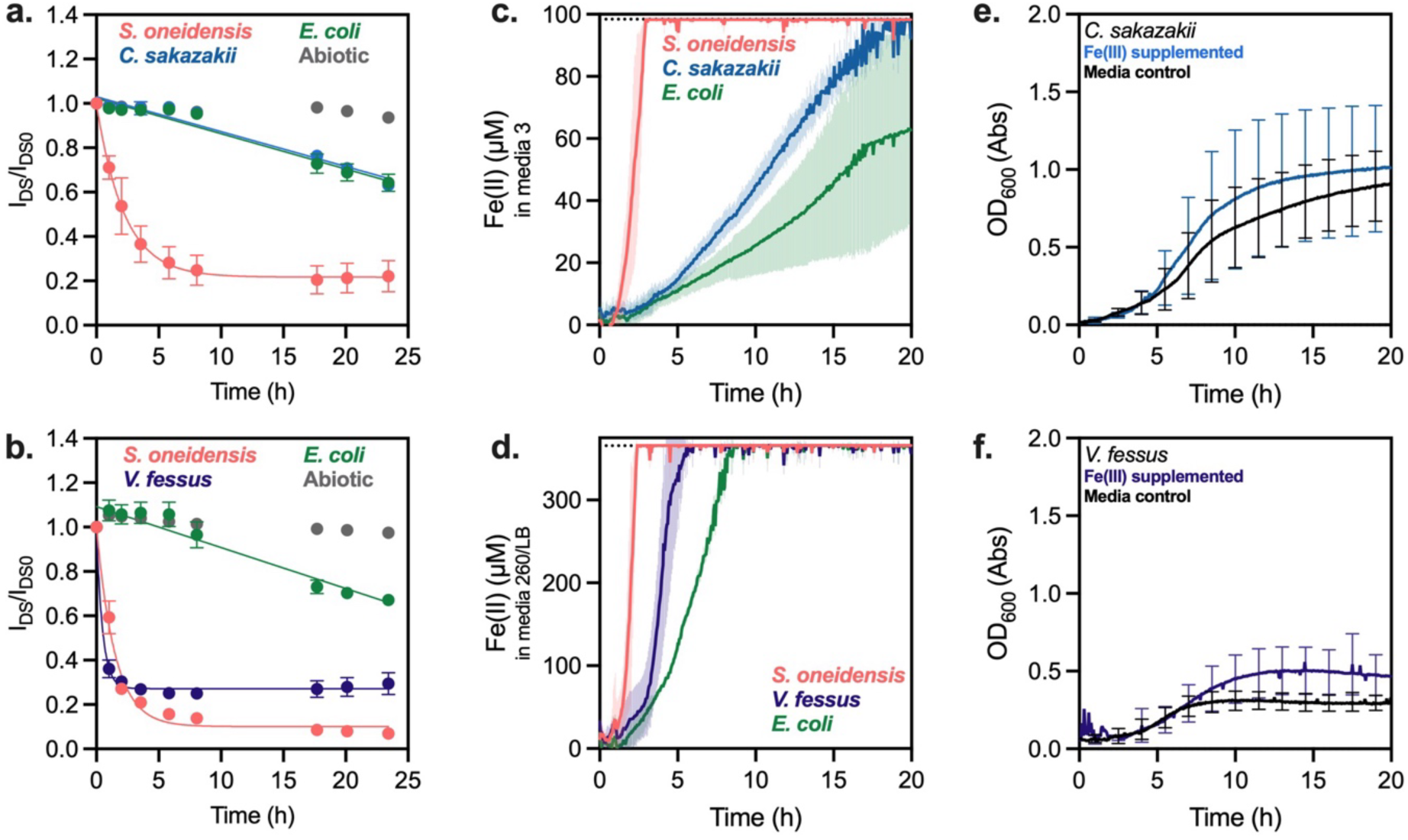
Examination of putative electrogens *C. sakazakii* and *V. fessus*. **a.-b.** Current over time from bacterially generated de-doping of a PEDOT:PSS-coated electrode. The I_DS_/I_DS0_ curve was prepared with an initial inoculum OD_600_=0.01. Curve for **a.** was collected in Media 3 for all species, and curve for **b.** was collected in a 1:50 mixture of Media 260:LB broth. **c.-d.** Fe(II) generation curves measured via ferrozine colorimetric assay over time. Initial inoculum of OD_600_=0.01 was from the same bacterial stock used to generate **a.** and **b.** respectively and run in the same respective media. Cells were supplemented with 5 mM Fe(III) at beginning of assay. **e.-f.** Growth curves for *C. sakazakii* and *V. fessus* with and without Fe(III) supplementation. Data represents n=3 ± standard deviation.

## Discussion

Microdroplet emulsions address key challenges of single-cell sorting when studying an extracellular process such as EET. Common methods for characterizing phenotypic differences within a population, such as FACS^80^, cannot be used for studying extracellular processes as signal cannot be attributed back to a source cell. In a microdroplet system, each cell is spatially isolated within its own pico- to nano- liter aqueous reaction vessel, allowing for smaller reaction sizes and higher sample numbers compared to a flask or even 96-well^81^ or 384-well plates traditionally used to study EET^79,82^. However, microdroplet systems require new, potentially unconventional assay development requirements^52^. A notable barrier was the requirement of a fluorescent signal that correlates to EET-activity. Our recent work linking Cu(I)-catalyzed cycloaddition to EET in *S. oneidensis* provided the opportunity to use a fluorescent probe for cycloaddition as an indirect measure for EET. Two further obstacles to success in developing a microdroplet screen for EET were cell survivability in droplet formation and oxygenation. The first was overcome by adapting a pico-injection protocol^46^, and visible growth within droplets was seen (Figure 2h and 2i). The second challenge, involving oxygenation, required a more unique solution. Many microdroplet system designed elements aim to introduce oxygen and oxygenation at multiple steps, including the oxygen-permeable fluorinated oil. To circumvent this problem, an oxygen-limited benchtop method was developed by degassing the starting solutions and plumbing the microfluidic system under a positive pressure (Figure 1, Figure S3, and Figure 2a-c). This created a closed system that could be maintained through collection into a nitrogen-filled syringe, and anaerobic bacteria could be grown in the droplet system. The ability to retrofit a bench-top system for oxygen-limited screens is advantageous in the study of anaerobic bacteria. This could prove to be useful in the study of anaerobic gut bacteria, of which some have been seen to be EET-capable, but their role is still poorly understood.

The amendments to the protocol allowed for a fluorescence output from CuAAC within *S. oneidensis* filled- droplets. CalFluor488^51^ is uniquely suited to the microdroplet system, as it undergoes a quenched to fluorescent conversion, allowing for low background in unreacted droplets. Additionally, the reaction is catalyzed by Cu(I), which can amplify signal and improve sensitivity as even a small amount of reduction can produce enough catalyst to yield fluorescent signal. While the droplet histograms are not diagnostic, populations can be sorted based on fluorescent output and sorting parameters were selected and recorded based on fluorescently gating the output. Utilizing this method, wild-type *S. oneidensis* and an EET-deficient knockout (Δ*mtrC*Δ*mtrF*Δ*omcA*) were mixed and screened for the ability to perform EET. We measured an approximately 2-fold enrichment of the wild-type in the sorted population, validating our ability to enrich for EET. The Δ*mtrC*Δ*mtrF*Δ*omcA* knockout is not a perfect negative control (Figure 2d), as it can still perform some reduction due to the remaining MtrA and MtrD proteins. Background reduction is something we and others have seen before^20,52,69,79,83^; however, our enrichment for MR-1 indicates that despite this background reduction we can capture electrogens in a mixed environment. Stricter sorts involving multiple sequential FADS could be used to isolate a more selective subset of bacterial species, where the sorted population would be collected and subjected to additional rounds of sorting. Microscopy images (Figure 2h and 2i) confirm that bacteria are growing within the microdroplets, and that within a mixed population a high fluorescent signal is not tied to solely cell growth but corresponds to the relative population density of the EET-capable phenotype.

Microfluidic systems are ideal for avoiding microbe-microbe competition for resources which can mute studies of their phenotypic behavior^47,49,80^. This is best illustrated by examining the prevenance of iron oxidation capable genomes in the lake water enrichment, which is increased in the bulk enrichment but decreased for the droplet system (Figure 4a). These data could be a result of the bulk enrichment where multiple species interact and affect the presence or absence of each other: as Fe(II) builds up due to the presence of iron reducers, there is an off-target advantage for iron-oxidizers which contain genes similar to MtoAB and Cyc2 (cluster 3) which have been implicated in iron oxidation^67^. If this is the case, it highlights a potential advantage of the microdroplet system as no genes from this cluster were identified from the example genomes in the droplet enriched population. When examining the genus distribution of the microdroplet screen of the lake water, encapsulation alone (prior to FADS) enriched notably for *Psuedomonas* (76.1% of the encapsulated population). These results are not surprising given that certain *Psuedomonas* species are known to secrete Cu-binding small molecules and the assay exposes the bacteria to a relatively high concentration of copper^84–86^. Consequently, there is likely a growth advantage in this particular genus due to co-encapsulation with Cu. Interestingly, despite the large percentage of *Psuedomonas* in the droplet population, no *Pseudomonas* was detected in any of the fluorescent sorted populations indicating that it was not actively catalyzing CuAAC despite its growth-based advantage. These results highlight the utility of the microfluidic system, despite the growth-based advantage observed with *Pseudomonas*, growth is separated from function in a way that cannot easily be done in traditional bulk EET-screens. Many traditional enrichments tie growth to function; however, our system allows the sorting to overcome the initial encapsulation bias. Consequently, the droplet system may be a powerful tool for identifying non-growth related EET, especially where mediated electron transfer may not necessarily be tied to the central carbon metabolism^25^.

We identified a number of interesting bacteria from droplet sorting of an environmental sample. Despite not previously being reported as EET-capable, several bacteria were identified by both bulk and droplet enrichment, indicating that they may have electrogenic behavior. *Acinetobacter lwoffii*, was originally isolated from an arsenic-polluted environment. It has been reported to co- exist with many heavy metal compounds, and was enriched in both droplet and bulk enrichments^87^. *Granulicatella adiacens*, is notoriously difficult to culture under laboratory conditions^88^, but has been reported in several cases of infection surrounding metallic hip replacements^89^. Our microdroplet screen identified a large increase in *G. adiacens*; however, it was present in a much-lower quantities in the bulk enrichment perhaps indicating its difficulty growing under typical laboratory conditions. Further, *Exiguobacterium indicum*, which has been shown to bio-reduce hexavalent chromium^90^, was similarly isolated by our screen and the bulk enrichment. *E. indicum* is also referenced as having the ability to reduce azo dyes^91^, an ability shared by *Shewanella* species and sometimes attributed to EET^92,93^. Additionally, known EET-capable organisms such as *Ochrobactrum* species, were enriched only in the microdroplet system where we had over 1000- fold enrichment^19^.

To validate species identified solely the microdroplet screen, we selected two species that were not previously reported as EET-capable, *C. sakazakii* and *V. fessus*, and analyzed them in monoculture. They both identified solely in the top 1.8% of performers (Sample V, tight gate), and we examined their ability to reduce Fe(III) and current. *C. sakazakii* is known to infect infant formula and has previously been the focus of study regarding its iron acquisition systems and ability for ferric iron to disrupt biofilm formation^76,77^, but the ability to generate current or perform EET has not been reported. To the best of our knowledge, there has not been a report of *V. fessus* displaying any iron-specific, heavy metal tolerance, or current generation. In monoculture, *V. fessus* was able to reduce the current at a rate comparable to that of *S. oneidensis*. Both bacteria generated Fe(II) from soluble Fe(III), although *C. sakazakii* only marginally outperformed the negative control of *E. coli*. Interestingly, it appears that neither bacteria, under our culturing conditions, was able to utilize Fe(III) as its sole terminal electron acceptor. Notably, we were unable to culture *V. fessus* without the presence of at least 0.1% sheep’s blood. Even when attempting to substitute with soluble Fe(III) citrate, the bacteria would not grow (Figure S7). These data in tandem with their identification in our microdroplet screen reinforces that we are able to look for non-growth-related electron transfer via microdroplets. We believe this indicates electrogenic behavior of these bacteria, particularly strong electrogenic behavior from *V. fessus* and potentially weak electrogenic behavior from *C. sakazakii*.

While there are some commonly accepted bacteria known to perform EET: *S. oneidensis*, *L. monocytogenes, G. sulfurreducens*; to the best of our knowledge, there is not a master list or completely agreed upon set of conditions for what may be considered an “EET-capable” or Fe(III)- reducing bacteria. While there are sets of genes and behaviors that indicate ability to perform EET, very few bacteria have been expressly tested for this phenotype and few agree what constitutes as an appropriate test for EET^11,12,34,94–96^. When compiling a literature search, including review articles, of EET-capable organisms existing lists often had little to no overlap between others. Of the articles we examined, the average number of species shared between the lists was 3 with standard deviation of 6 species^26,31,37,44,63–66^, indicating that we needed to additional tools for examining our data. For this reason, we utilized the FeGenie program^67^, which looks at potential iron-related genes and their relative abundance within a set of genomes as an indirect method for looking for EET-ability. This program was designed to be used with full genome sequencing not with 16S data. As a result, example genomes were gathered from the NIH National Center for Biotechnology Information (NCBI), and we were limited to only to the representative genomes available. These example genomes most likely do not fully capture the information potential in the data set or the power of the FeGenie program. For that reason, we limited identification to the presence of one or more genes in each pathway as opposed to genomic frequency of homologous genes. Of the 366 starting bacteria, full genome sequencing was only available for 222 genomes through NCBI with many being members of a specific genus but not of a known species name. This highlights the power of partnering a phenotype screen with full genome sequencing that could fully utilize the capabilities of FeGenie. However, the diverse nature of electrogens and lack of consensus emphasizes that genotyping alone is incomplete as a mechanism of identifying EET-capable bacteria. This underscores the necessity of a phenotype screen to further inform the genotype-phenotype relationship.

The potential difference selective pressures between the droplet and bulk enrichment indicates that while this microdroplet screen is useful particularly at identifying *S. oneidensis*-like EET, for which it was developed, it should serve to complement traditional EET screening such as growth on electrodes^31,45,97,98^, dialysis enrichments^60,99,100^, growth on iron^34,35,101^, and more. This is likely due to a few reasons: the requirement for survival in the emulsion, the relatively high concentration of Cu within the droplets, and the use of Cu(II) rather than Fe(III) for an extracellular electron acceptor. Notably, current methods for identifying EET activity tend to rely on growth-related behavior, such as the ability to use Fe(III) as a sole terminal electron acceptor, or the ability to outcompete other bacteria within a bulk enrichment or in biofilm ^8,29,102–104^. There exists a need to study both weak electrogens, and non-growth associated EET for their complex role in the environment, mineral-microbe interactions, and microbe-microbe interactions^25,30,44,105^. Furthermore, some bacteria exhibit EET only under certain conditions which are not always captured *in vitro*, such as during pathogenesis^106^ or under specific substrate and potential conditions^107^. Further work focused on increasing cell viability and detecting Fe(III) reduction as opposed to Cu(II) reduction would further aid in investigating electrogens in complex environments. In summary, this work represents a step towards high-throughput methods for identifying specific types of EET and for isolating bacteria based on phenotype in a microdroplet emulsion.

## Materials and Methods

CalFluor 488 (Click Chemistry Tools), alkyne-PEG4-acid (Click Chemistry Tools), copper(II) bromide (CuBr_2_, Sigma-Aldrich, 99%), 2-(4-((bis((1-(tert-butyl)-1H-1,2,3-triazol-4- yl)methyl)amino)methyl)-1H-1,2,3-triazol-1-yl)acetic acid (BTTAA, Click Chemistry Tools >95%), sodium DL-lactate (NaC_3_H_5_O_3_, TCI, 60% in water), sodium fumarate (Na_2_C_4_H_2_O_4_,VWR, 98%), EPES buffer solution (C_8_H_18_N_2_O_4_S, VWR, 1 M in water, pH = 7.3), potassium phosphate dibasic (K_2_HPO_4_, Sigma-Aldrich), potassium phosphate monobasic (KH_2_PO_4_, Sigma-Aldrich), sodium chloride (NaCl, VWR), dimethyl sulfoxide (cell culture grade, Sigma-Aldrich), ammonium sulfate ((NH_4_)_2_SO_4_, Fisher Scientific), magnesium(II) sulfate heptahydrate (MgSO_4_·7H_2_O, VWR), trace mineral supplement (ATCC), casamino acids (VWR), Lysogeny Broth (BD), OptiPrep (Sigma-Aldrich), Pico-Gen^TM^ 60 x 60 single aqueous (Sphere Fluidics), Pico-Wave ^TM^ (Sphere Fluidics), Pico-Break ^TM^ 1 (Sphere Fluidics), Pico-Mix ^TM^ (Sphere Fluidics), Pico-Surf^TM^ (Sphere Fluidics), Medical Grade Polyethylene Micro Tubing / 0.015" ID x 0.043" OD (+/- .003") = .38mm ID x 1.09mm OD (+/- .076mm) / (100’ Roll) (Scientific Commodities), *Cronobacter sakazaki* (Farmer et al.) Iverson et al. (ATCC 29544), *Vagococcus fessus* Hoyles et al. (ATCC BAA-289), defibrillated sheep’s blood (Lampire), Tryptic Soy Broth (BD 211825), Nutrient Broth (BD cat 234000), Agar (BD), disodium;4-[3-pyridin-2-yl-6-(4- sulfonatophenyl)-1,2,4-triazin-5-yl]benzenesulfonate hydrate (Ferrozine, VWR) All media components were autoclaved or sterilized using 0.2 μm PES filters.

### Oxygen-Limited Encapsulation of *S. oneidensis*

Overnight cultures were grown in a Coy Anaerobic Glovebox containing a humidified atmosphere at 3% hydrogen content and the balance nitrogen. The cultures were started by picking a single colony into argon-sparged LB broth supplemented with 20 mM sodium lactate (2.85 µL of 60% w/w sodium lactate per 1 mL culture). After overnight growth anaerobically at 30°C, cell cultures were diluted to an OD_600_ of 0.00006 into a solution of 40 mM lactate, 80 mM fumarate, 20 wt% OptiPrep in LB broth. Inside of the anaerobic chamber, 1 mL of the cell solution was loaded into the aqueous syringe (BD). The tubing was prepared by heat-sealing one end, and the other was loaded onto a needle, inside of the anaerobic glove box, the needle was attached to an empty syringe and pressure was pulled three times and held for 30 seconds each time and then vented after each round to remove any O_2_ present in the tubing. Additionally, 10 mL of argon-sparged 2.5% 008-FluoroSurfactant in Pico-Wave^TM^ was loaded into the oil syringe (SGE) and capped with a sealed, sparged needle and tubing. A collection syringe was prepared by pulling 6 mL of the Coy Anaerobic Glovebox atmosphere and capped with a sealed, sparged needle and tubing. A PicoDroplet Single Cell Encapsulation System (Sphere Fluidics) was used. All sealed syringes were removed from the anaerobic chamber. Each syringe was loaded under a 100 µL/h positive pressure before clipping the heat-sealed end and connecting to the 60 × 60 droplet maker (Pico- Gen) chip. To make the emulsion, the syringe pumps were increased to 1000 µL/h (aqueous) and 1200 µL/h (oil). The system was allowed to calibrate before cutting and plumbing the collection syringe (BD). After encapsulation, emulsions were incubated at 30 °C 24 h to allow for growth within the droplets.

### Encapsulation of Mixed population

Overnight cultures were grown in a Coy Anaerobic Glovebox containing a humidified atmosphere at 3% hydrogen content and the balance nitrogen. The cultures were either started by picking a single colony into argon-sparged LB broth supplemented with 20 mM sodium lactate (2.85 µL of 60% w/w sodium lactate per 1 mL culture) or 20 mM dextrose (for *E. coli* Nissle 1917 and *S. cerevisiae* BY4741). After overnight growth anaerobically at 30°C, cell cultures were diluted to an OD_600_ of 1.8 x 10^5^ (MR-1), 7.8 x 10^5^ (EcN), and 0.008 (*S. cer*) into a solution of 40 mM lactate, 40 mM glucose, 80 mM fumarate, 20 wt% OptiPrep in LB broth. These reflect a loading density overall of 1 in 10 droplets, and the loading ration of 30:35:35 (MR-1:EcN:*S. cer*). The remaining encapsulation is the same as that of *S. oneidensis* outlined above.

### Oxygen-Limited Pico-Injection of CuAAC components

A 5X Cu(I)-catalyzed Alkyne—Azide Cycloaddition solution (70 µM CalFluor 488, 2 mM Alkyne-Peg4-acid, 2 mM Cu:BTTAA (1:6)) was prepared from a 3.2 mM stock of CalFluor 488 in DMSO, 8 mM stock of CuBr_2_ in water, 48 mM stock of BTTAA in water and a 4 mM stock of Alkyne-Peg4-Acid in water in a Coy Anaerobic Glovebox containing a humidified atmosphere at 3% hydrogen content and the balance nitrogen. Into a 1 mL syringe, 200 µL of the solution was loaded into the aqueous syringe (BD) and capped with a sealed, sparged needle and tubing. Additionally, 10 mL of argon-sparged Pico-Wave^TM^ was loaded into the oil syringe (SGE) and capped with a sealed, sparged needle and tubing. A collection syringe (12 mL Manufacterer) was prepared by pulling 6 mL of the Coy Anaerobic Glovebox atmosphere, adding in 500 µL of 008- Fluorosurfactant (5%) in Pico-Wave^TM^, and capped with a sealed, sparged needle and tubing. The droplets were then transferred to a 1 mL syringe (BD) and capped with a sealed, sparged needle and tubing. All sealed syringes were removed from the anaerobic chamber, loaded onto the syringe pumps set at 15 µL/h (aqueous), 100 µL/h (droplets) and 1200 µL/h (oil) before cutting off the sealed end and plumbing into a Pico-Mix^TM^ chip. The system was allowed to calibrate before cutting and plumbing the collection syringe (PGE). After picoinjection, emulsions were incubated at 30 °C 24 h to allow for growth within the droplets.

### Fluorescent Sorting for heterogeneity

For droplet sorting, a Single Cell Assay and Isolation platform (Sphere Fluidics) with a 488 nm laser 244 and 525/50 emission filter (GFP) was used. PMT setup of the system was set to a gain of 0.8 or 1 such that an appropriate spread was seen without maxing out the detector for both PMT 1 (GFP channel/bandpass 525/50) and PMT 2 (large bandpass 650/150). Peak detection minimum was set at 0.07 and maximum at 100. The minimum width was set to 0.18 and maximum set to 100. Sorting gates were determined based on population distribution and manually drawn above 1 V(MR-1), 2.1V, 3.5V (Lake water), 4.25V (Lake water enrichment round 2). The top 3000 droplets were then collected using an applied voltage of 0.3V. After sorting, the emulsion was broken with 100 μL Pico-Break (Sphere Fluidics) and with extracted or cultured. Culturing occurred in LB broth supplemented with 20 mM lactate at 30 °C overnight. Each sample was split, freezing 500 µL as a cryogenic stock, and diluting 10 µL into the anaerobic chamber to start the creation of the next generation of sort.

### Lake water enrichment

Environmental and post-droplet enriched samples were enriched in an anaerobic bulk system microcosom similar to those previously described^61^. Briefly, 10 mL of sediment containing lake water containing a sediment-associated mixed microbial community was inoculated into a 250- mL anaerobic bulk systems filled with filter-sterilized lake water. The cells were provided with lactate at 0.1mM as a carbon source, and iron-containing sediments were isolated in 3.5 kDa dialysis tubing. The cells were allowed to grow for 5 days and redox potential, pH, and aqueous Fe(II) concentrations were measured daily.

### 16S Sample Extraction and sequencing

DNA samples from both pre- and post-enriched samples via the oxygen-limited droplet protocol and bulk sedimentation enrichments were purified via the AllPrep DNA/RNA kit from Qiagen per manufacturer’s instructions. Samples were sent to Mr. DNA for microbial taxonomic analysis. Mr. DNA utilizes 515F-806R (V4 region) primers, and classification is determined using QIIME^108^. Taxonomic classification from Mr. DNA was utilized at the genus and species level for all subsequent analysis.

Species richness for each sample was estimated using a rarefaction curve based on species level counts. The rarefaction curve data was generated using the ‘rarecurve’ function in the vegan (v. 2.6.4)^109^ library with default settings. Taxonomic proportions were calculated by dividing counts from each individual taxa by the total number of read counts for that sample. We estimated a species to show prospective ‘enrichment’ for downstream testing within a sample based on the following criteria: if a certain taxa was above 0.015% of the initial population, and also showed a higher proportion in the compared population of interest. We estimated this initial population detection cutoff based on the initial library size and the consistency of the number of enrichments for a specified sample. Taxa of interest were also selected if they showed notable non-zero detection within samples of interest, while showing no observance in reference samples. Species enrichments were used to obtain a list of genomes to later be used for FeGenie. Analyses were performed using custom scripts in python and R. Taxonomic barplots and rarefaction plot were generated using ggplot2 (v. 3.5.0)^110^. The EET species database was generated via Genomic sequences from these taxa were collected for any full genome or 16S (full or V4) sequences available on NCBI from the following citations^18,19,26,31,44,63–65,65,66,105,111^. Plots were generated in R.

### FeGenie

Example genomes were obtained using NCBI and chosen based on the longest whole genome sequencing available for a given genome. If the species level was not specified, the most appropriate example genome was chosen from the genus. Each genome was exported and FeGenie was run on Python3 and the code was available for download on GitHub. Results were then processed to normalize the number of genes per genome analyzed and represented as percentages. These graphs were created using Python3 and the code is available in the SI. Relative fold-enrichment was calculated by normalizing the enriched population by the starting population.

### OECT device operation and electrochemistry

Two different types of electrolytes were used according to the bacterial strains. Medium 3 was used with *Cronobacter sakazakii*, *S. oneidensis* MR-1, and *E. coli*. A mixture of medium 260 and Luria-Bertani (LB) broth at 1:50 was used with *Vagococcus fessus*, *S. oneidensis*, and *E. coli*. All electrolytes were purged with argon bubbling for at least 15 minutes before use. Before each experiment, the OECT slides and PDMS layer were autoclaved separately and subsequently assembled in the biosafety cabinet. The fabrication of devices is outlined further in the Supplementary Information. To ensure an oxygen-free environment, the OECT experiments were carried out inside the nitrogen-filled glovebox. Electrochemical measurements were conducted with the multichannel potentiostat (MultiPalmSens4, PalmSens BV). Prior to inoculation, OECTs were equilibrated in the glovebox with abiotic electrolytes for 3 days. For the inoculation process of the OECTs, all cells were grown anaerobically overnight in their respective media. Grown cell cultures cell density (OD_600_) measured, the cells were concentrated and resuspended to the intended OD600 at 0.1 with the respective purged media, forming the inoculum culture. Subsequently, the inoculum cultures were used to inoculate OECTs at a 1:9 ratio of cell culture to the OECT electrolyte, achieving an intended inoculation OD_600_ at 0.01. During the OECT experiment, the gate voltage VGS and drain voltage VDS were constantly biased at 0.2 V and -0.05 V, respectively. Transfer curve measurements, were conducted 24 hours after inoculation, with the V_DS_ kept at-0.05 V while the V_GS_ scanned from -0.1 V to 0.6 V with a scan rate of 20mV/s. The open circuit potentials (OCP) of the source and gate electrode potentials were measured against the Ag/AgCl pellet reference electrodes (RE) (550010, A-M Systems). The Ag/AgCl electrodes were directly inserted into the OECT chamber without using any salt bridges.

### Ferrozine Assay

Aerobic cultures of *S. oneidensis, E. coli*, *C. sakazakii*, and *V. fessus* were created in each bacteria’s preferred rich media (LB, LB, Media 3 and Media 260 respectively). The next day, cultures were diluted 1/100 into anaerobic media and allowed to grow overnight. Each bacteria was washed via centrifugation at 6000 rcf and decanted before the supernatant was exchanged for the reaction media (Media 3 for *C. sakazakii* and Media 260:LB 1:50 for *V. fessus*). Cultures of *S. oneidensis* and *E. coli* were also washed and reconstituted in each media to be run concurrently. Ferrozine was dissolved into degassed, anaerobic growth media such that the final concentration of the assay can be run at 1 mg/mL. 6.57 µL of 190 mM Fe(III)citrate was added in addition to 10 µL of cells directly from aerobic overnights into a 250 µL reaction in a clear-bottom U shaped Grenier 96- well plate (final Fe(III) concentration of 2 mM). A calibration curve of Fe(II) sulfate stocks in sterile water were inoculated into control wells for each media containing ferrozine from 12 µM- 0 µM. Each media had a cell-free control inoculated with Fe(III) but no cells. The reduction of Fe(II) was measured using the calibration curve minus the background reduction via the media blank. The complete 96-well plate was sealed anaerobically and placed into an incubated plate reader. Readings were taken every 3 minutes at 562 nm. The mixed cultures of Δ*mtrC*Δ*mtrF*Δ*omcA* and wild-type MR-1 were grown overnight in *Shewanella* Basal Media (SBM) supplemented with 0.05 % casamino acids and Wolfe’s Mineral Solution. The cells were used without washing and subjected to the same treatment in SBM with casamino acids and mineral solution.

### Growth on Fe(III)

To a 96-well plate, 6.57 µL of 190 mM Fe(III)citrate was added in addition to 10 µL of cells directly from aerobic overnights into a 250 µL reaction in a clear-bottom U shaped Grenier 96- well plate in the anaerobic growth media (Media 3 for *C. sakazakii,* and 1:50 Media 260:LB for *V. fessus*). Fe(III)-free wells were made with the addition of water in the place of Fe(III). Media controls were included for blanking the readings. The complete 96-well plate was sealed anaerobically and placed into an incubated plate reader. Readings were taken every 3 minutes at 600 nm.

## Supporting information

Droplets_SI

## Acknowledgements

Collections of genomes were downloaded from the NIH with assistance by Kenneth C. Sabjel and Pedro Sobral. *S. oneidensis* Δ*mtrC*Δ*omcA*Δ*mtrF* and ϕ,Mtr were generously provided by Prof. Jeffrey Gralnick (U. Minnesota). The photo of Town Lake in Figure 3 was taken by Emma J Palmer. This research was financially supported by the Welch Foundation (Grant F-1929, B.K.K.), the National Institutes of Health under award number R35GM133640 (B.K.K.), an NSF CAREER award (1944334, B.K.K.), and the Air Force Office of Scientific Research under award number FA9550-20-1-0088 (B.K.K.). G.P. and E.J.P were supported through National Science Foundation Graduate Research Fellowships (Program Award No. DGE-1610403). The Sphere Fluidics system was funded by a Cooperative Agreement (W911NF-17-2-0091) between Army Research Laboratory (ARL) and UT Austin. Opinions, conclusions, interpretations, and recommendations are those of the authors and are not necessarily endorsed by the US Army. The mention of trade names or commercial products does not constitute endorsement or recommendation for use by the Department of the Army or the Department of Defense. Schematics were created using BioRender.com, graphs were created in Prism GraphPad, R, and Python. 16S data was analyzed using R.

## Author Contributions

G.P., E.K.B., H.S.A., and B.K.K. conceived the project and research design. G.P. and E.B. designed and performed microdroplet emulsion experiments. G.P. ran CuAAC control experiments and analyzed putative electrogens. E.J.P. ran bulk enrichment and extracted DNA for sequencing. Y.G. ran OECT experiments and characterized responses. R.R. and G.P analyzed 16S sequencing results. H.S.A. and B.K.K. supervised research. G.P., H.S.A., and B.K.K wrote the manuscript with input from all authors.

## Materials & Correspondence

Request for materials and correspondence should be addressed to H.S.A. and B.K.K.

## Data Availability

Experimental data supporting the findings of this study will be available through the Texas Data Repository.

