## Supplementary material for "Single-Cell Phenotyping of Extracellular Electron Transfer via Microdroplet Encapsulation": Droplets_SI

Supplementary Figure 1 Fluorescent conversion of CalFluor488 to fluorogenic triazole. ........ 5

Supplementary Figure 2. Chemical emulsion controls of CuAAC on microdroplet system ........ 5

Supplementary Figure 3. Schematic for oxygen-limited microdroplet emulsion generation, pico-injection, and sorting ........ 6

Supplementary Figure 4. Histogram of FADS sorting of a mixture of S. oneidensis MR-1, E. coli Nissle 1917, and S. cerevisiae BY4741. ........ 6

Supplementary Figure 5. Image of 96-well plate after ferrozine assay for Fe(II) detection. ........ 7

Supplementary Figure 6. Rarefaction curves for 16S data gathered from differing enrichment conditions. ........ 8

Supplementary Figure 7. Anaerobic growth of putative electrogen*s* ........ 9

Supplementary Figure 8. Transfer curves 24 hours post inoculation ........ 9

***Code Availability*** ........ 10

R-code for 16S proportionality charts (Figure 3) ........ 10

Python Code for FeGenie Dot Plot ........ 11

### Supplementary Methods

#### CuAAC Chemistry Controls

Fluorescent assay controls were completed in 96-well plate format outlined previously[1]. Briefly, fluorescence emission was collected on a BMG LABTECH CLARIOstar plate reader with a 491 (±14) nm and an emission collection at 538 (±38) nm). Different to previously published methods, all reactions were performed in LB broth supplemented with lactate (20 mM) as a carbon source and fumarate (20 mM) as the primary electron acceptor. Stock solutions of 1 M sodium fumarate and 60 w/v% lactate solutions were stored at 4 °C until use. Aliquots of 3 mM and 11 mM of CalFluor 488 were created in DMSO and stored frozen at −80 °C until use. Aliquots of 4 mM alkyne-PEG$_4$-acid were created in sterile water and stored at −20 °C until use. An 8 mM copper bromide stock in sterile water was created and stored at 4 °C and mixed with an equal volume amount of 48 mM freshly made stock of BTTAA in sterile water. Control reactions included increasing the concentration of CalFluor 488, running with new buffer conditions, and using a dual phase system where both Pico-SURF ™ and Pico-WAVE ™ were added in 100 µL volumes to the 96-well plate and the LB-reaction mixture was run in a 100 µL volume floating on top of the oil layer. The reaction was then placed into the plate reader for analysis and allowed to react for between 10 and 24 h.

#### Fe(III)-reduction detection for Δ*mtrC*Δ*omcA*Δ*mtrF* and MR-1 mixed cultures

After running the Ferrozine assay, the endpoint absorbance at 562 nm was collected. The average and standard deviation of 9 samples of MR-1 was used to determine the "ON" conditions. If the absorbance of a given sample was within one standard deviation of the MR-1 values, it was bucketed into MR-1, and otherwise it was considered a knockout. Visual plate layout and image in Figure S5.

#### Culturing conditions and notes for non-model organisms

The bacteria *C. sakazakii* and *V. fessus* are non-model organisms and were cultured in their preferred media as described on ATCC (Media 3 and 260 respectively). When culturing *V. fessus*, we found that at least 0.1% of the media must be sheep's blood to support growth. The organism appeared to grow to comparable densities in Media 260 and in Media 260 that had been diluted in LB up to 50 X. However, we found for optimal growth and health, the bacteria were best cultured in Media 260 liquid after inoculation from either 260 agar plate or glycerol (22%) stock before being diluted into other media conditions. *C. sakazakii* could be grown in both LB and Media 3 and seemed more robust to changes in its culturing conditions. *V. fessus* could be grown with or without the presence of 5% $CO_2$ at 37°C shaking at 250 rpm. However, *C. sakazakii* grew best under non-shaking conditions at 30°C, and experienced growth defect when grown with shaking at 200 or 250 rpm. Both bacteria could be grown anaerobically at 30°C without shaking and in their preferred media reached approximately and $OD_{600}$ of 0.2 and 0.7 for *C. sakazakii* and *V. fessus* respectively (Figure SX). Both bacteria were unable to grow anaerobically in M9Y media but *C. sakazakii* saw a significant improvement in growth when supplemented with 1 mM Fe(III) as an electron acceptor.

#### Device Fabrication

Fabrication of the organic electrochemical transistors (OECTs) was guided by methodologies outlined in prior work[2]. The quartz microscope slides (AdValue Technology, Model FQ-S-003) were cleaned with a sequence of soapy water, acetone, followed by isopropyl alcohol, and dried using nitrogen gas. The slides were then subjected to oxygen plasma treatment using reactive-ion etching technology (RIE) at an output of 150 W and oxygen flow rate of 50 sccm for 120 seconds. Post-etching, the slides were layered with AZ5209E photoresist and went through photolithography to set the electrode layout. After the patterning process, quartz substrates received a deposit of 10 nm titanium (Ti) adhesion layer and subsequent 100 nm gold (Au) electrode layer through thermal evaporation. Residual materials were then cleared away utilizing the acetone lift-off approach. A second photolithographic procedure was executed to delineate the PEDOT:PSS area covering both the transistor channel and gate-tip, achieving dimensions of 150 µm by 10 µm and 500 µm by 500 µm, respectively. The PEDOT:PSS solution (CleviosTM PH1000) was filtered through a 0.22 µm PES filter, followed by deposition onto the prepared quartz slides using a spin-coating technique. The coated slides were then placed on a hot plate set at 90 °C for 15 minutes to dry, and the process was completed with an acetone lift-off. To enhance the electrical conductivity, the PEDOT:PSS films were heated on a hot plate at 90 °C and submerged in ethylene glycol for 3 minutes. The OECT chamber layers comprised of casted polydimethylsiloxane (PDMS, SylgardTM 184) mixed with a 9% weight proportion of curing agent. The mixture was carefully drop-cast onto the mode and left to solidify over 48 hours at ambient temperature to ensure an even surface finish. The OECT chambers and the fluid access ports in the PDMS layers were cut out with hole punches.

**Supplementary Table 1.** Bacterial strains and plasmids used in this study.

| Strain or plasmid | Description/Genotype | Reference or source |
|---|---|---|
| *S. oneidensis* Strains MR-1 | MR-1 (ATCC700550), wild-type strain accession code NC_004347 | American-Type Culture Collection |
| *S. oneidensis* Δ*mtrC*Δ*omcA*Δ*mtrF* | Lacks outer membrane cytochromes MtrC, OmcA, and MtrF; Δ*mtrC*Δ*omcA*Δ*mtrF* (JG596) | Jeffrey Gralnick, U. of Minnesota |
| *S. oneidensis* ΔMtr-pathway | Lacks all outer membrane cytochromes (JG1194) | Jeffrey Gralnick, U. of Minnesota |
| *E. coli* Nissle 1917 | Wild-type strain accession code CP007799 | MetaFluor Probiotics |
| *E. coli* K12 MG1655 | Wild-type strain U00096 | Lydia Contreras, U. of Texas at Austin |
| *C. sakazakii* (strain CDC 4562-70) | *Cronobacter sakazakii* (Farmer et al.) Iversen et al. (ATCC 29544) accession code CP011047 | ATCC 29544 |
| *V. fessus* | *Vagococcus fessus* Hoyles et al. (ATCC BAA-289) accession code AJ243326 | ATCC BAA-289 |
| *S. cerevisiae* BY4741 | *Saccharomyces cerevisiae* Meyen ex E.C. Hansen (ATCC 201388) accession code 4022422 | ATCC 201388 |

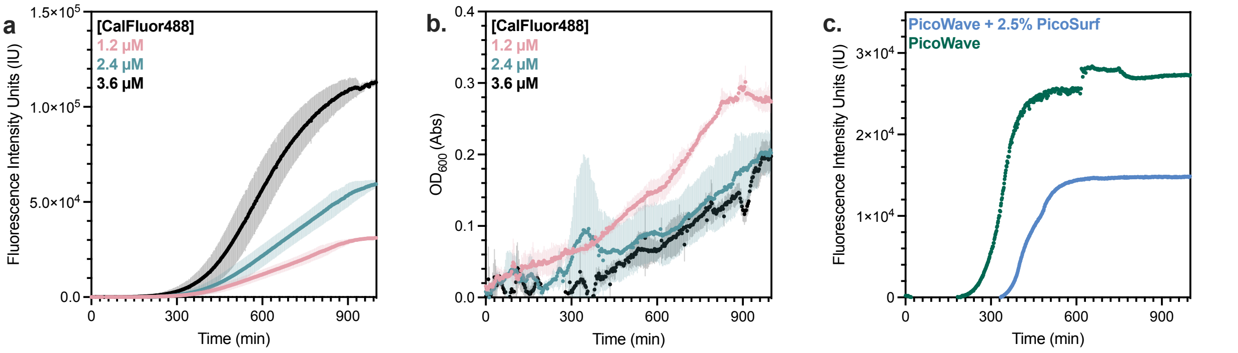

**Figure S1. Fluorescent conversion of CalFluor488 to fluorogenic triazole**. **a.** *S. oneidensis* MR-1 catalyzed CuAAC in the presence of increasing equivalents of reactants. **b.** Growth as measured by $OD_{600}$ for the reactions in **a.** over the course of their reaction. Data represent the mean of n=3 biological triplicates ± standard deviation. **c.** CuAAC reaction catalyzed by *S. oneidensis* MR-1 in the presence of fluorinated oil (PicoWave) and surfactant (PicoSurf). Data is in singlicate.

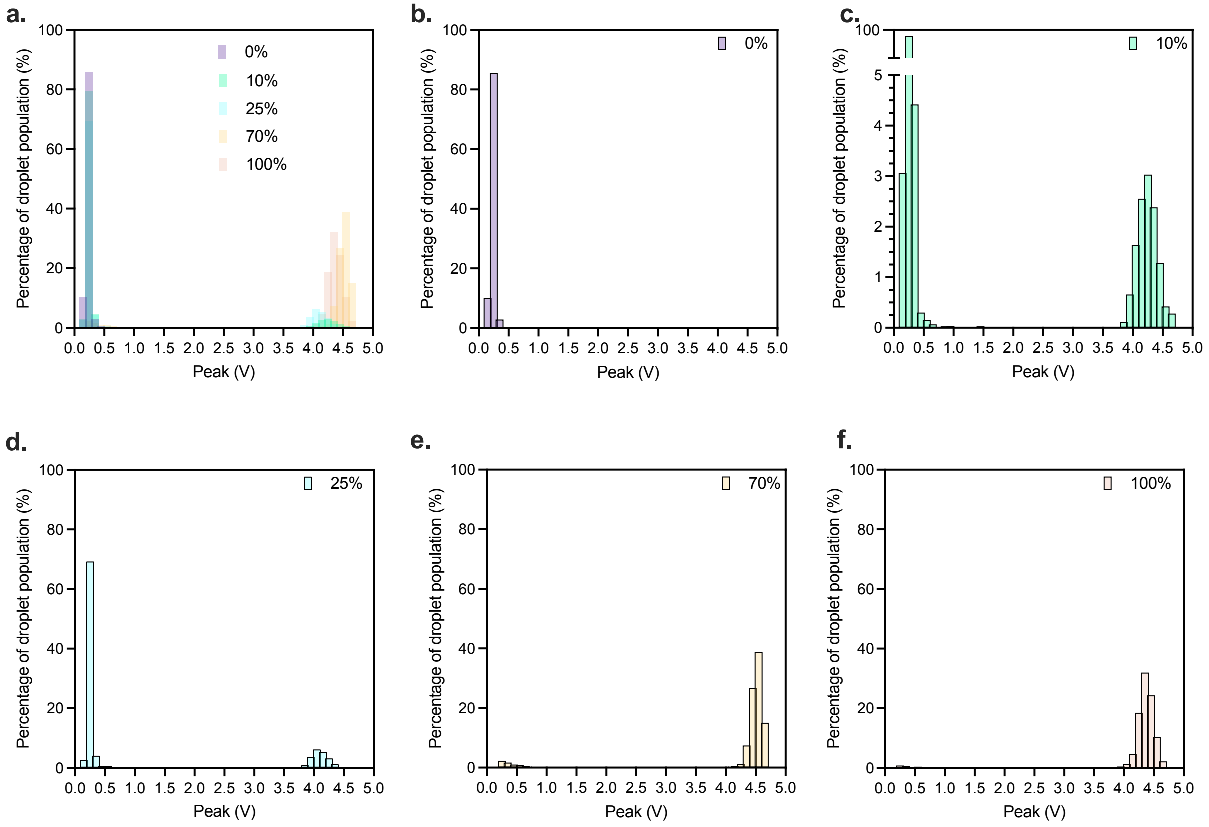

**Figure S2**. **Chemical emulsion controls of CuAAC on microdroplet system**. **a.** Summary data of all chemically reacted emulsions. Percentages indicate the amount of fully reacted emulsion mixed with unreacted starting material. **b.-f.** histograms with varying percentages of reacted emulsion. The y-axis of **c.** has been changed to allow for better viewing of reacted population.

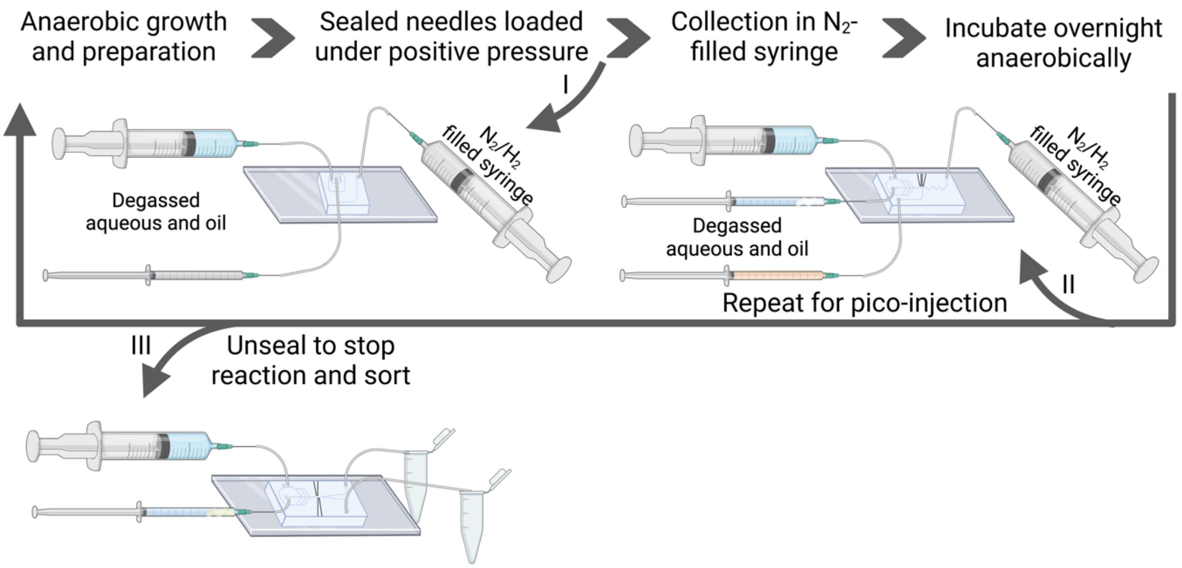

**Figure S3. Schematic for oxygen-limited microdroplet emulsion generation, pico-injection, and sorting.** Anaerobic growth and preparation of solutions for oxygen-limited emulsion.

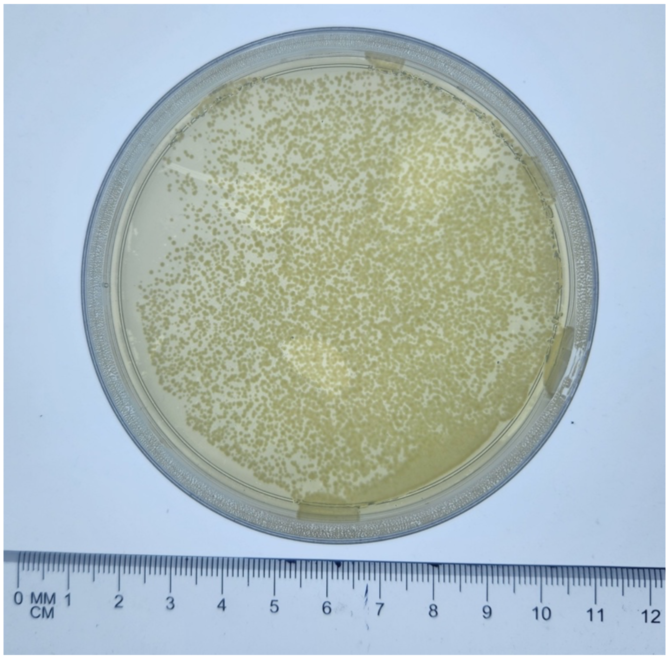

**Figure S4. *S. oneidensis* ΔMtr-Pathway post-sort recovery.** Emulsion collected from Figure 2c, broken, and plated. Plate was allowed to grow for 16h at 30°C before being imaged.

a. b.

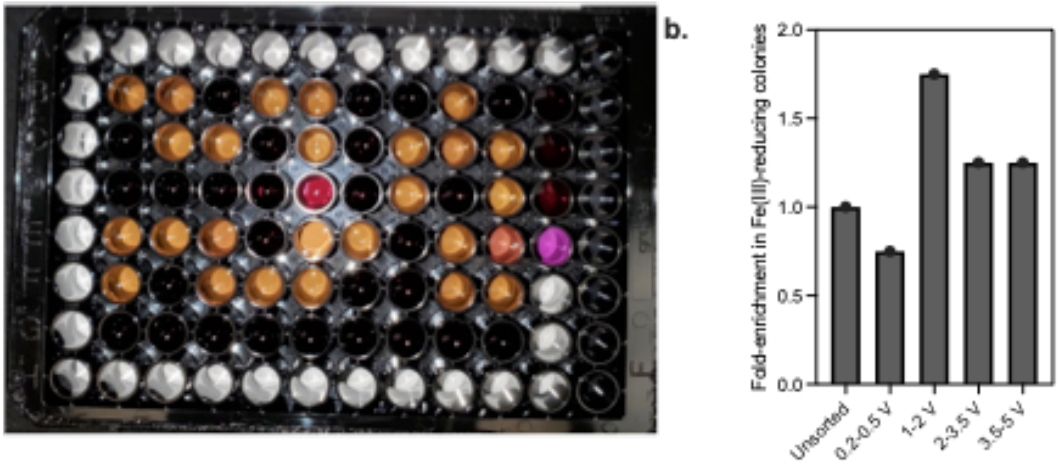

c.

| | 2 | 2 | 3 | 4 | 5 | 6 | 7 | 8 | 9 | 10 | 11 |
|---|---|---|---|---|---|---|---|---|---|---|---|
| B | Unsorted | Unsorted | Unsorted | Unsorted | Unsorted | Unsorted | Unsorted | Unsorted | Unsorted | Unsorted | Ferrozine Controls |
| C | 0.2-0.5V | 0.2-0.5V | 0.2-0.5V | 0.2-0.5V | 0.2-0.5V | 0.2-0.5V | 0.2-0.5V | 0.2-0.5V | 0.2-0.5V | 0.2-0.5V | Ferrozine Controls |
| D | 1-2V | 1-2V | 1-2V | 1-2V | 1-2V | 1-2V | 1-2V | 1-2V | 1-2V | 1-2V | Ferrozine Controls |
| E | 2-3.5V | 2-3.5V | 2-3.5V | 2-3.5V | 2-3.5V | 2-3.5V | 2-3.5V | 2-3.5V | 2-3.5V | 2-3.5V | |
| F | 3.5-5V | 3.5-5V | 3.5-5V | 3.5-5V | 3.5-5V | 3.5-5V | 3.5-5V | 3.5-5V | 3.5-5V | 3.5-5V | |
| G | MR-1 Control | MR-1 Control | MR-1 Control | MR-1 Control | MR-1 Control | MR-1 Control | MR-1 Control | MR-1 Control | MR-1 Control | MR-1 Control | |

**Figure S5. Image of 96-well plate after ferrozine assay for Fe(II) detection**. **a.** Image of 96-well plate post assay with outline of configuration **(c.)** of the plate. Samples from this were collected from FADS in Figure 2f. **b.** Fold-enrichment of iron reducing colonies as seen in figure **a.**

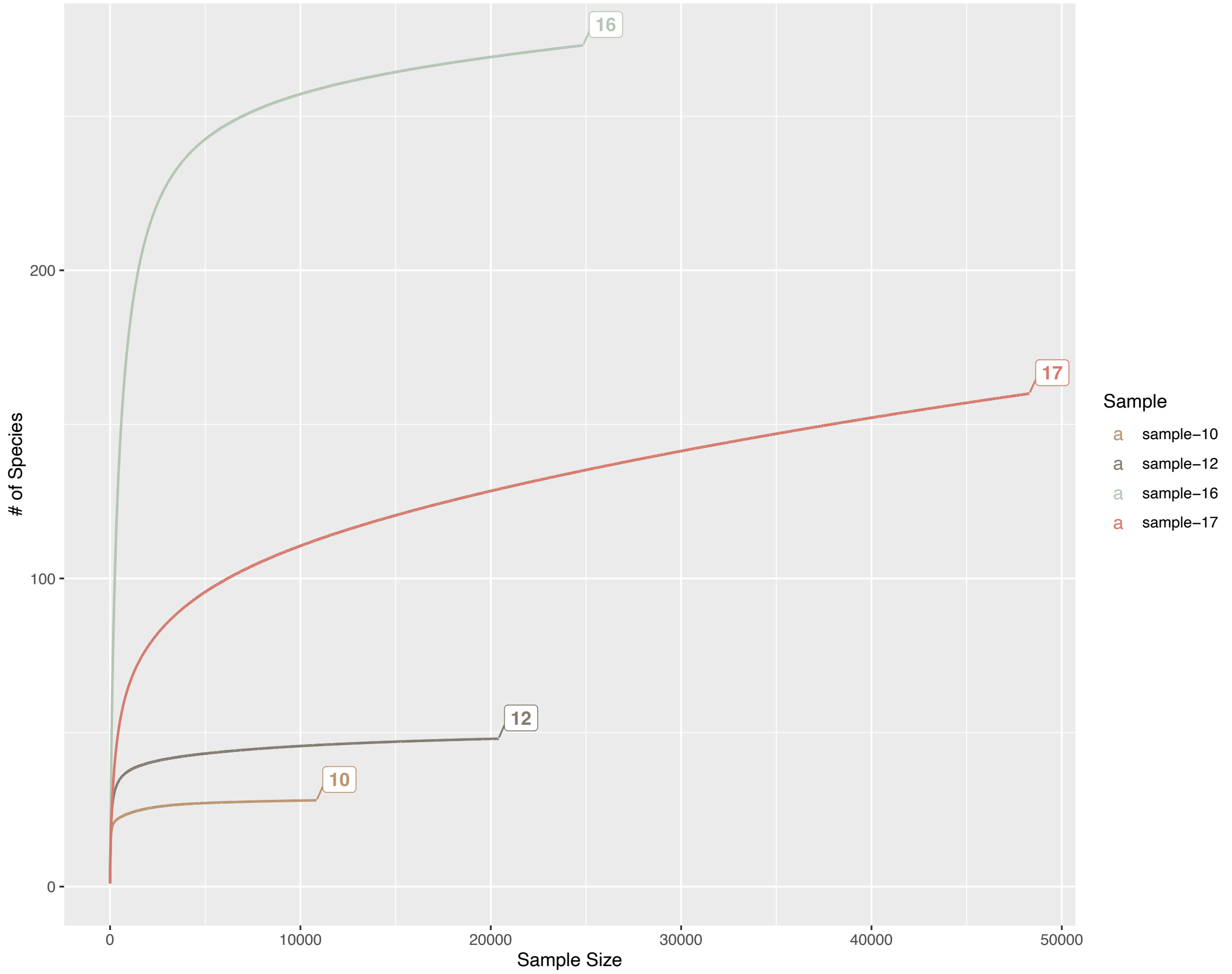

**Figure S6. Rarefaction curves for 16S data gathered from differing enrichment conditions**. Rarefaction curves generated for sample of reads from 16S sequencing. Sample-16 is the starting material from Lake Austin that was split. Sample-17 is the bulk enrichment on Fe-sedimentation. Sample-10 is the wide gated (over 2.5V) droplet enrichment, and sample-12 is the tight gate (over 3.1V) droplet enrichment.

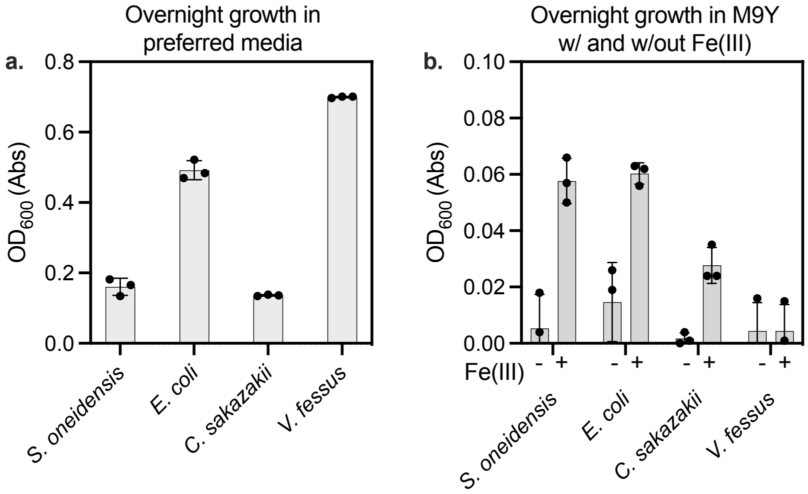

**Figure S7. Anaerobic growth of putative electrogens**. **a.** Overnight anaerobic growth in preferred media for *S. oneidensis* (*Shewanella* basal media with 0.05wt% casamino acids and Wolfe's mineral solution), *E. coli* K12 MG1655 (Lysogeny Broth), *C. sakazakii* (ATCC Media 3), *V. fessus* (ATCC Media 260). **b.** Overnight anaerobic growth in minimal M9Y media with and without Fe(III) supplementation. Data represents the mean of *n*=3 biological replicates with error bars ± the standard deviation.

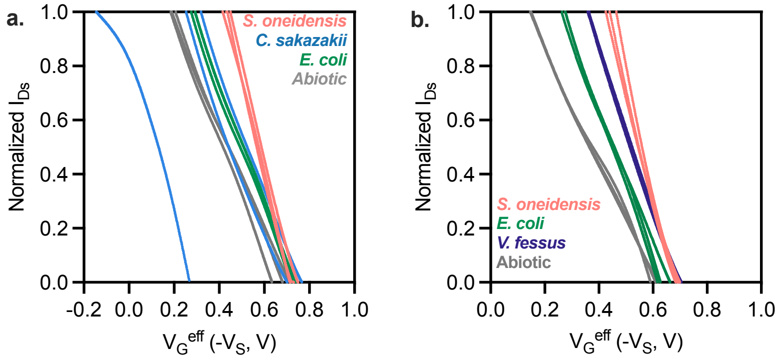

**Figure S8. Transfer curves 24 hours post inoculation**. **a.** Transfer curves plotted against the effective gate voltage in ATCC media 3 after 24 hours. **b.** Transfer curves plotted against the effective gate voltage in ATCC media 260 diluted 50X into Lysogeny Broth (LB).

**Python and R-Code**

**R-Code for 16S proportionality charts (Figure 3).**

```
library(readxl)
library(tidyverse)
library(data.table)
library(ggfortify)
library(plotly)
library(paletteer)
library(grid)
library(ggrepel)
library(gridExtra)
library(dplyr)
library(ggplot2)
library(devtools)
library(RColorBrewer)

# Set working directory
setwd("/Users/fig3_data/")

# Load data from RDS file
vals1_df <- readRDS("/Users/fig3_data/vals1_df.rds")

## Figure 3
positions = c('sample-16','sample-17',
        'sample-9','sample-10','sample-12')
cols_df_fig1 = tibble(taxa = sort(unique(vals1_df$taxa)), colors =cols_fig1)
cf1 =  deframe(cols_df_fig1)

cols_fig1 = colorRampPalette(paletteer_d("palettetown::pelipper",direction = 1))(98)
cols_fig1[[1]] = "#B49080FF"
fig1_barplot = ggplot(data = vals1_df, aes(x=sample,y= freq,fill=taxa)) +
  geom_bar(stat = "identity",width=0.8) +
  scale_fill_manual(values = cf1) +
  scale_x_discrete(limits = positions,
          position = 'bottom',
          labels=c('I.','II.','III.','IV.', 'V.')) +
  ggtitle("") + theme_bw() +
  theme(
    axis.text.x = element_text(angle = 90, vjust = 0, hjust = 1, face = "bold", size = 7),
    axis.text.y = element_text(angle = 90, hjust = 0.5, size = 7, color = "black"),  # Set y-axis label color to
black
    axis.title.y = element_text(size = 7, face = "bold", color = "black"),  # Set y-axis title color to black
    legend.position = "none",
    panel.border = element_rect(colour = "black", fill = NA, size = 1),  # Set panel border color to black
    panel.grid.major = element_blank(),  # Remove major gridlines
    panel.grid.minor = element_blank()   # Remove minor gridlines
  ) +
  ylab("Proportion") +
  xlab("")
fig1_barplot
```

```
## Making legend
vtop_g1 = vals1_df %>% group_by(taxa) %>% summarise(all = sum(freq)) %>% arrange(desc(all))
legend_top_f1 = vtop_g1[1:20,]
legend_top_f1$sample = "S1"
fig1_leg = ggplot(data = legend_top_f1, aes(x=sample,y= all,fill=taxa)) +
  geom_bar(stat='identity') + scale_fill_manual(values = cf1,name="Genus")

legend <- cowplot::get_legend(fig1_leg)
grid.newpage()
grid.draw(legend)
```

**Python code for FeGenie Dot-Plot**

```python
import pandas as pd

import numpy as np
import matplotlib.pyplot as plt
import matplotlib.colors as mcolors
from matplotlib.cm import ScalarMappable
import textwrap  # Import the textwrap module

# Read the first Excel file for color mapping
df_color = pd.read_excel('PercentPresent.xlsx')

# Read the second Excel file for dot sizes
df_size = pd.read_excel('presentfoldenrich.xlsx')

# Extract the x-axis values from the first row
x_values = df_color.columns[1:]

# Extract the y-axis values from column A and reverse the order
y_values = df_color.iloc[:, 0][::-1]

# Extract the matrix of values for the heatmap and reverse the order of
rows
matrix_color = df_color.iloc[:, 1:].values[::-1]
matrix_size = df_size.iloc[:, 1:].values[::-1]  # New matrix for dot sizes

# Create a figure and axis for the heatmap
fig, ax = plt.subplots(figsize=(12, 6))  # Adjust the figure size as
desired

# Define a custom color map for the heatmap
```

```
colors = [(0.8, 0.8, 0.8), (0, 0.6, 0), (0, 0.6, 0.6), (0, 0.6, 0.6), (0,
0.1, 0.6), (0, 0, 0.4)]  # Light Grey, Green, Dark Blue
cmap_heatmap =
mcolors.LinearSegmentedColormap.from_list('custom_colormap', colors,
N=256)

# Plot the heatmap with transparent squares
heatmap = ax.imshow(matrix_color, cmap=cmap_heatmap, alpha=0, extent=[-
0.5, len(x_values) - 0.4, -0.5, len(y_values) - 0.4])

# Non-linear scaling factor for circle size
nonlinear_scale_factor = 0.15
# Add circles to represent the values with colors based on the custom
colormap and sizes from the new file
for i in range(len(y_values)):
    for j in range(len(x_values)):
        value_color = matrix_color[i, j]
        value_size = matrix_size[i, j]
        radius = value_size * nonlinear_scale_factor  # Apply square root
scaling to circle size
        color = cmap_heatmap(heatmap.norm(value_color))  # Use the custom
colormap for circles
        circle = plt.Circle((j, i), radius=radius, color=color, alpha=0.7)
        ax.add_patch(circle)

# Draw a new rectangle that surrounds only the data from the Excel file
(excluding the last row)
rect = plt.Rectangle((-0.5, -0.4), len(x_values) + 0.22, len(y_values) -
0.1, edgecolor='black', linewidth=1, facecolor='none')
ax.add_patch(rect)

# Use textwrap to wrap labels at spaces and slashes with earlier wrapping
ax.set_xticks(np.arange(len(x_values)))
ax.set_yticks(np.arange(len(y_values)))
ax.set_xticklabels([textwrap.fill(label, width=15, break_long_words=False)
for label in x_values])
ax.set_yticklabels([textwrap.fill(label, width=15, break_long_words=False)
for label in y_values])

# Rotate the x-axis tick labels if needed

# Add a legend with circles representing values from the new Excel file
legend_values = [0, 0.5, 1.5, 2.5, 3.5]  # Adjusted legend values based on
the new size mapping
```

```
legend_x_offset = len(x_values) + 0.3  # Adjusted legend offset to prevent
cutoff
legend_y_offset = -1.1  # Adjust vertical position of the legend

for i, value in enumerate(legend_values):
    scaled_value = value * nonlinear_scale_factor  # Apply square root
scaling to legend circles
    circle = plt.Circle((legend_x_offset, i + legend_y_offset),
radius=scaled_value, color='gray', alpha=0.7)
    ax.add_patch(circle)
    ax.text(legend_x_offset + 1, i + legend_y_offset, f"{value}",
color='black', ha='left', va='center')

ax.set_xlim(-0.5, len(x_values) + 2.5)
ax.set_ylim(-0.5, len(y_values) - 0.5)  # Adjusted ylim to start from -0.5

# Remove the black rectangle surrounding the entire graph
ax.set_frame_on(False)

# Calculate the actual minimum and maximum values from your data
data_min = np.min(matrix_color)
data_max = np.max(matrix_color)

# Add a new legend with a colorbar
sm = ScalarMappable(cmap=cmap_heatmap, norm=plt.Normalize(vmin=data_min,
vmax=data_max))
sm.set_array([])  # You need to set a dummy array for the ScalarMappable
cax = fig.add_axes([0.035, 0.17, 0.02, 0.8])  # Adjusted legend position
cbar = plt.colorbar(sm, cax=cax, orientation='vertical', label='  % of
genomes containing one or more genes related to:', shrink=0.3, pad=0.0)  #
Adjust the shrink and pad as needed
cbar.ax.yaxis.set_label_position('left')  # Set label position to the left
cbar.ax.tick_params(axis='y', direction='inout')  # Set tick direction to
both inside and outside

# Adjust the figure layout to include the title and legend outside the
plot area
plt.tight_layout()

plt.savefig('heatplotdotplot.jpg', format='jpg', dpi=250)

# Show the plot
plt.show()
```
